# Axin-mediated regulation of lifespan and muscle health in *C. elegans* involves AMPK-FOXO signaling

**DOI:** 10.1101/2020.04.22.055962

**Authors:** Avijit Mallick, Ayush Ranawade, Bhagwati P Gupta

## Abstract

Aging is a significant risk factor for several diseases. Studies have uncovered multiple signaling pathways that modulate the process of aging including the Insulin/IGF-1 signaling (IIS). In *C. elegans* the key regulator of IIS is DAF-16/FOXO whose activity is regulated by phosphorylation. A major kinase involved in DAF-16-mediated lifespan extension is the AMPK catalytic subunit homolog, AAK-2. In this study, we demonstrate a novel role of PRY-1/Axin in AAK-2 activation to regulate DAF-16 function. The *pry-1* transcriptome contains many genes associated with aging and muscle function. Consistent with this, *pry-1* is strongly expressed in muscles and muscle-specific overexpression of *pry-1* extends the lifespan, delays muscle aging, and improves mitochondrial morphology in DAF-16-dependent manner. Furthermore, PRY-1 is necessary for AAK-2 phosphorylation. Together, our data demonstrate a crucial role of PRY-1 in maintaining the lifespan and muscle health. Since muscle health declines with age, our study offers new possibilities to manipulate Axin function to delay muscle aging and improve lifespan.

## INTRODUCTION

Aging is defined as a progressive functional decline in living organisms. It is characterized by hallmarks such as genomic instability, epigenetic alterations, mitochondrial dysfunction, and telomere attrition, and is thought to be regulated in part by genetic pathways (López-Otín et al., 2013). Although this process of aging had generated immense curiosity, it was only in the early 1980s that the researchers started to investigate aging in the nematode *Caenorhabditis elegans* (Klass, 1983). Since then, several genetic factors and pathways have been reported to govern and modulate lifespan that is conserved in higher eukaryotes (Lapierre and Hansen, 2012; Uno and Nishida, 2016). Insulin/insulin-like growth factor-1 signaling (IIS) was the first pathway shown to be involved in the regulation of aging in *C. elegans* (Kenyon, 2011; Kenyon et al., 1993). Subsequent studies showed that the IIS pathway is conserved across eukaryotes (Uno and Nishida, 2016). In *C. elegans* reduction in IIS receptor homolog, DAF-2, activity leads to a prolonged lifespan that is dependent on DAF-16, a FOXO transcription factor homolog (Kenyon et al., 1993). This modulation of lifespan by DAF-16 involves translocation to the nucleus followed by either the activation or repression of genes involved in stress response, metabolism and autophagy (Lee et al., 2003; Meléndez et al., 2003; Murphy et al., 2003).

The activity of DAF-16 is regulated by phosphorylation (Lin et al., 2001). One of the kinases involved in this process is the α2 catalytic subunit homolog of AMPK, AAK-2 (Greer et al 2007a), a phenomenon that appears to be conserved in the mammalian system (Greer et al. 2007b). AAK-2 also plays a crucial role in aging. It is essential for DAF-2-mediated lifespan elongation and its overexpression extends the lifespan of animals (Apfeld et al., 2004). Interestingly, a truncated version of AAK-2, bearing only the catalytic domain, was found to be more effective than the full-length wild-type form, suggesting that AAK-2 activity is regulated during the normal aging process (Mair et al., 2011). Similar to the *C. elegans*, AMPK in *Drosophila* is also involved in lifespan regulation. Specifically, overexpression of the α2 subunit in muscles and fat bodies extended the lifespan of transgenic animals (Stenesen et al., 2013).

AMPK is an established energy sensor in eukaryotes that is phosphorylated by several kinases including LKB1 (Burkewitz et al., 2014; Hardie et al., 2012). Studies in the mouse and human cell culture models have shown that under the condition of glucose starvation AMPK forms a complex with LKB1 and the scaffolding protein, Axin (Zhang et al., 2013b), The multimeric complex regulates AMPK activation, leading to phosphorylation of downstream targets (Hardie and Lin, 2019; Hardie et al., 2012). The involvement of Axin in AMPK complex formation is essential since Axin knockdown drastically reduces AMPK activity, leading to fatty liver in starved mice (Zhang et al., 2013b). In addition to their role in AMPK regulation, Axin family members are also involved in multiple biological processes during development and post-development (Mallick et al., 2019a). Since its discovery as a negative regulator of WNT signaling, Axin is demonstrated to participate in other, non-WNT, pathways as well. In all cases, a common thread is Axin’s role as a scaffold protein to recruit other factors to form complexes (Mallick et al., 2019a). However, whether the scaffolding role of Axin affects FOXO activity remains to be investigated.

In this study, we report that the *C. elegans* Axin homolog PRY-1, which is necessary for the embryonic and larval processes, is also essential for the normal lifespan maintenance. Previously, metformin-mediated lifespan extension was shown to depend on another *C. elegans* Axin-like gene, *axl-1*, however, *axl-1* does not play a role in aging and age-related processes (Chen et al., 2017). We found that animals lacking *pry-1* function during adulthood had a shorter lifespan, which was due to deterioration in associated processes as judged by the analysis of age-linked markers. Consistent with this, *pry-1* transcriptome contains aging-related miRNA and protein-coding genes. *pry-1* is broadly expressed in adults with high levels in body wall muscles (BWM). We also found that muscle-specific knock-down of *pry-1* caused an increase in the proportion of fragmented mitochondria and led to a reduction in lifespan. Conversely, overexpression of *pry-1* in muscles improved both these phenotypes. Thus, *pry-1* appears to affect lifespan by regulating muscle mitochondria health. Interestingly, muscle-specific expression of mouse Axin (*mAxin1*) in *C. elegans* also extended the lifespan of animals, suggesting that Axin’s role in aging may be conserved. It is worth mentioning that Axin is known to be expressed in mouse and human skeletal muscles (Smith et al., 2019; Uhlén et al., 2015). Our molecular genetic experiments revealed that PRY-1’s role in aging depends on AAK-2 and DAF-16. Moreover, we found that PRY-1 is necessary for AAK-2 phosphorylation, which in turn promotes nuclear localization of DAF-16 in the intestine. Taken together, these results allow us to suggest that PRY-1 interacts with AAK-2, presumably by forming a complex, leading to AAK-2 phosphorylation and regulation of DAF-16 function during lifespan maintenance.

Our work provides the first evidence of an in vivo role of Axin family member in promoting longevity and muscle health. Studies in humans and other higher systems have established a connection between aging, muscle health, mitochondrial dysfunction, and diseases (Gouspillou and Hepple, 2016; Hood et al., 2019). Furthermore, Axin is essential for muscle maintenance since myogenesis is abrogated in mutant animals (Huraskin et al., 2016) and *Axin2* upregulation is associated with increased muscle fibrosis in the aging mice (Arthur and Cooley, 2012; Brack et al., 2007). Since muscle mass and function progressively decline with age, understanding the mechanism of Axin’s function in this tissue promises to uncover potential interventions for age-associated muscle deterioration.

## RESULTS

### *pry-1* transcriptome contains genes involved in lifespan regulation

The involvement of PRY-1 in multiple signaling pathways and biological events is well documented (Mallick et al., 2019a). Earlier, we reported both mRNA and microRNA (miRNA) transcriptome profiles of *pry-1* mutant that revealed 2,665 differentially expressed (DE) protein-coding genes and six miRNAs (Mallick et al., 2019b; Ranawade et al., 2018). The characterization of DE genes revealed *pry-1*’s role in miRNA-mediated seam cell development (Mallick et al., 2019b) and its novel function in lipid metabolism (Ranawade et al., 2018). In this study, we have specifically focused on the genes linked to aging. The analysis of the mRNA transcriptome dataset shows that aging-related protein-coding genes and gene families are overrepresented (69 in total, 33% upregulated and 67% downregulated) (**Figure 1A; Table S1**). Of the DE miRNAs, *mir-246* is known to be involved in aging and stress response (De Lencastre et al., 2010).

**Figure 1:**
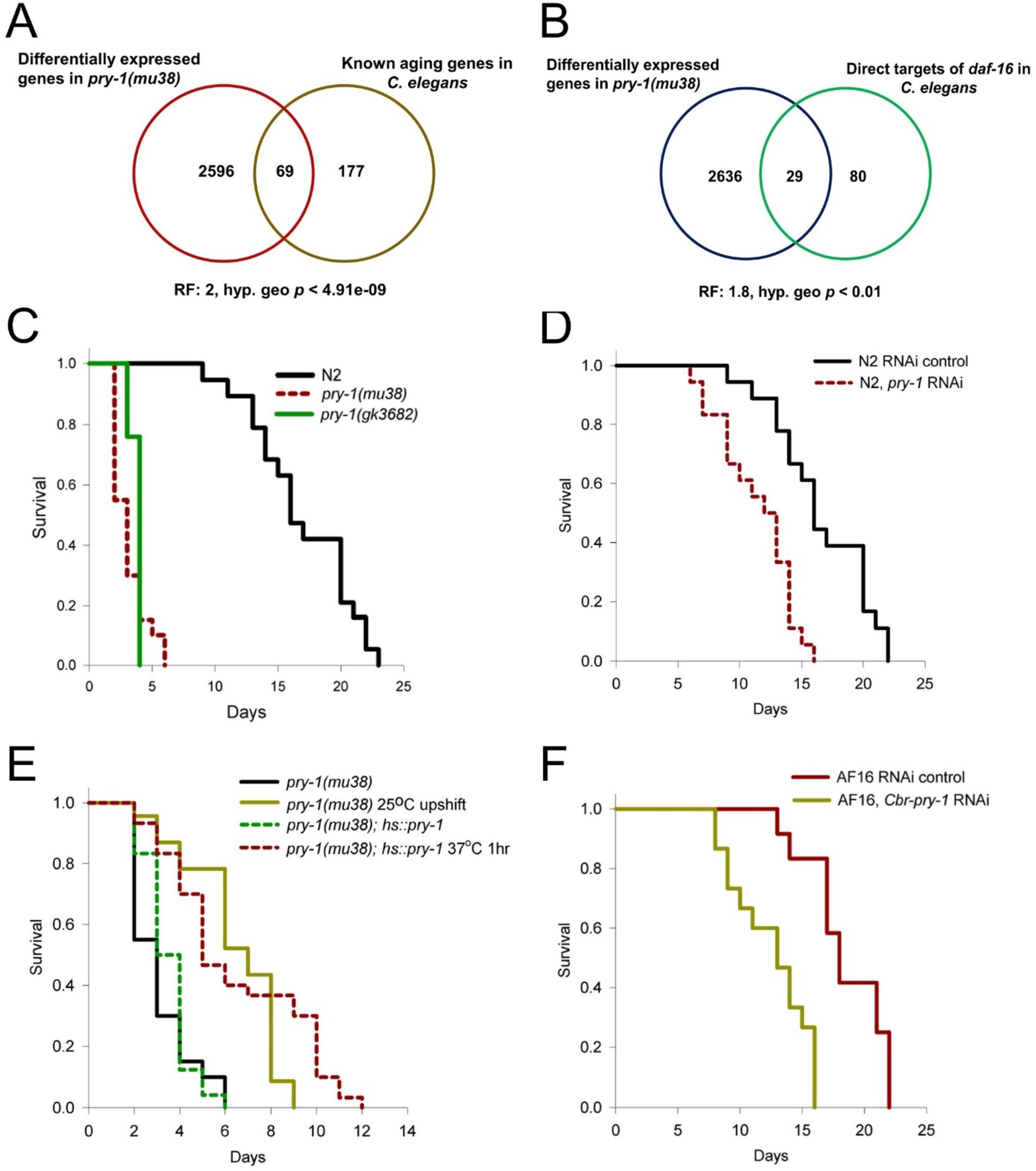
*pry-1* is required for the normal lifespan of animals. (A) 69 DE genes in *pry-1(mu38)* transcriptome are linked to aging. (B) 29 of the DAF-16 direct targets are present in *pry-1* transcriptome. (C) Mutation in *pry-1* results in reduced lifespan. (D) *pry-1* knockdown in *C. elegans* adults leads to shorter lifespan. (E) lifespan defect of *pry-1(mu38)* animals is rescued after temperature upshift to 25°C and ectopic *pry-1* expression using *hs∷pry-1* during adulthood. (F) Like *C. elegans pry-1*, RNAi of *Cbr-pry-1* causes animals to die early. For all the lifespan data with statistics see also STAR methods and Table S2.

GO term analysis was used to further analyze the 69 protein-coding DE genes, which identified associated biological activities such as cellular processes (26 genes), metabolic processes (24 genes) and biological regulation (13 genes). Within the cellular processes, candidates include genes linked to lipid metabolism (*aap-1, hyl-1, elo-2*, *ctl-2, cat-1,* and *lipl-4*), which further demonstrate the essential role of lipids in *pry-1*-mediated signaling (Ranawade et al., 2018), and suggest that *pry-1* may affect lipid metabolism to regulate aging.

The aging-related genes are linked to 32 distinct signaling pathways and include well-known factors such as AAP-1 (PI3K adapter subunit) and DAF-16, both belonging to the IIS pathway (Lapierre and Hansen, 2012, Uno and Nishida, 2016), and XBP-1, (human XBP1 ortholog) that acts downstream of IRE-1 and PEK-1-mediated signaling (Ron and Walter, 2007). Thus, *pry-1* appears to interact with multiple genetic networks. We also compared the *pry-1* transcriptome with DE genes of the DAF-2-DAF-16 signaling pathway (Lin et al., 2018) and found a significant overlap (415 genes; R.F.:1.7, *p*< 2.228e-31, **Table S3**). Additionally, 29 of 109 DAF-16 direct targets (Li and Zhang, 2016) are present in the *pry-1* dataset (27% overlap; two-thirds downregulated) including 4 (*dod-17, prdx-3, nnt-1,* and *daf-16*) that are directly involved in aging (**Figure 1B; Table S3**). Taken together, these *in silico* analyses suggest that *pry-1* acts in part via DAF-16 to regulate lifespan in *C. elegans*.

### Mutations in *pry-1* reduce the lifespan

We followed up the above results by performing experiments. The examination of animals carrying a nonsense mutation in *pry-1*, *mu38*, revealed an 80% reduction in the mean lifespan (**Figure 1C; Table S2**). A similar phenotype is also observed in a CRISPR allele, *gk3682,* that deletes roughly 750 bp region including the 5’ UTR and first exon (Mallick et al., 2019b) (**Figure 1C; Table S2**). As *pry-1* is also involved in developmental processes (Mallick et al., 2019a), we took an RNAi approach to knock down the gene function specifically during adulthood. As expected, *pry-1(RNAi)* animals were found to be short-lived, with a 22-31% reduction in the mean lifespan (**Figure 1D; Table S2**). Consistent with these results, *pry-1* transcript was significantly higher in older adults **(Figure S1A)**.

To further examine *pry-1*’s role in aging, we performed two additional experiments. In one case the cold-sensitive allele *mu38* was used. While the lifespan defect of *pry-1(mu38)* is severe at 20°C (mean lifespan 81% lower than N2, **Figure 1E; Table S2**), the animals appear healthier and show an improved lifespan at 25°C (50% lower than N2, **Figure 1E; Table S2**). When day-1 *pry-1(mu38)* adults were upshifted from 20°C to 25°C, lifespan defect was suppressed by two-folds (6.4 +/− 0.4 days mean lifespan compared to 3.2 +/− 0.1 days for untreated *mu38* control). The second experiment involved generation of transgenic animals carrying a heat-shock promoter-driven *pry-1.* The *hs∷pry-1* transgene efficiently rescued the lifespan defect of *pry-1(mu38)* animals upon heat-shock to day-1 adults (58% longer mean lifespan, **Table S2; Figure 1E**). Interestingly, no such effect was observed in wild-type animals **(Figure S1C)**.

Since our lab had previously demonstrated a role for *pry-1* ortholog in *C. briggsae* vulva development (Seetharaman et al., 2010), we also investigated *Cbr-pry-1’s* role in aging. The results revealed that similar to *C. elegans*, a reduction in the function of *C. briggsae pry-1* exhibited lifespan defects. Specifically, both mutation (*sy5353*) and RNAi-mediated knockdown of *Cbr-pry-1* during adulthood caused animals to age faster (**Figure 1F and S1B; Table S2**). These data demonstrate a conserved role of *pry-1* in lifespan maintenance in nematodes.

### *pry-1* knockdown in adults causes accelerated aging and increased expression of stress response markers

Several physiological and molecular changes occur in animals during the aging process. These include a decline in tissue function, oxidative stress, accumulation of mis/unfolded proteins, and altered lipid distributions (Huang et al., 2004; López-Otín et al., 2013). To characterize such changes in *pry-1(RNAi)* animals, we analyzed the age-dependent decline in pharyngeal pumping and body movement. Adult-specific knockdown of *pry-1* led to a significant reduction in pharyngeal pumping starting day-7 of adulthood (**Figure 2A**). Likewise, body movement analysis showed defects wherein the rate of body bends progressively decreased over the life of animals, beginning day-2 of the adulthood (**Figure 2B**). Together, these results demonstrate a pattern of accelerated physical deterioration in animals due to reduced *pry-1* activity. Similar phenotypes were also observed in *pry-1(mu38)* mutants although they were more severe (**Figure S2A and S2B**). Consistent with the adult-specific role of *pry-1*, we found that heat shocked *pry-1(mu38); hs∷pry-1* adults showed significant improvements in both body bending and pharyngeal pumping phenotypes (**Figure S2C and S2D**).

**Figure 2:**
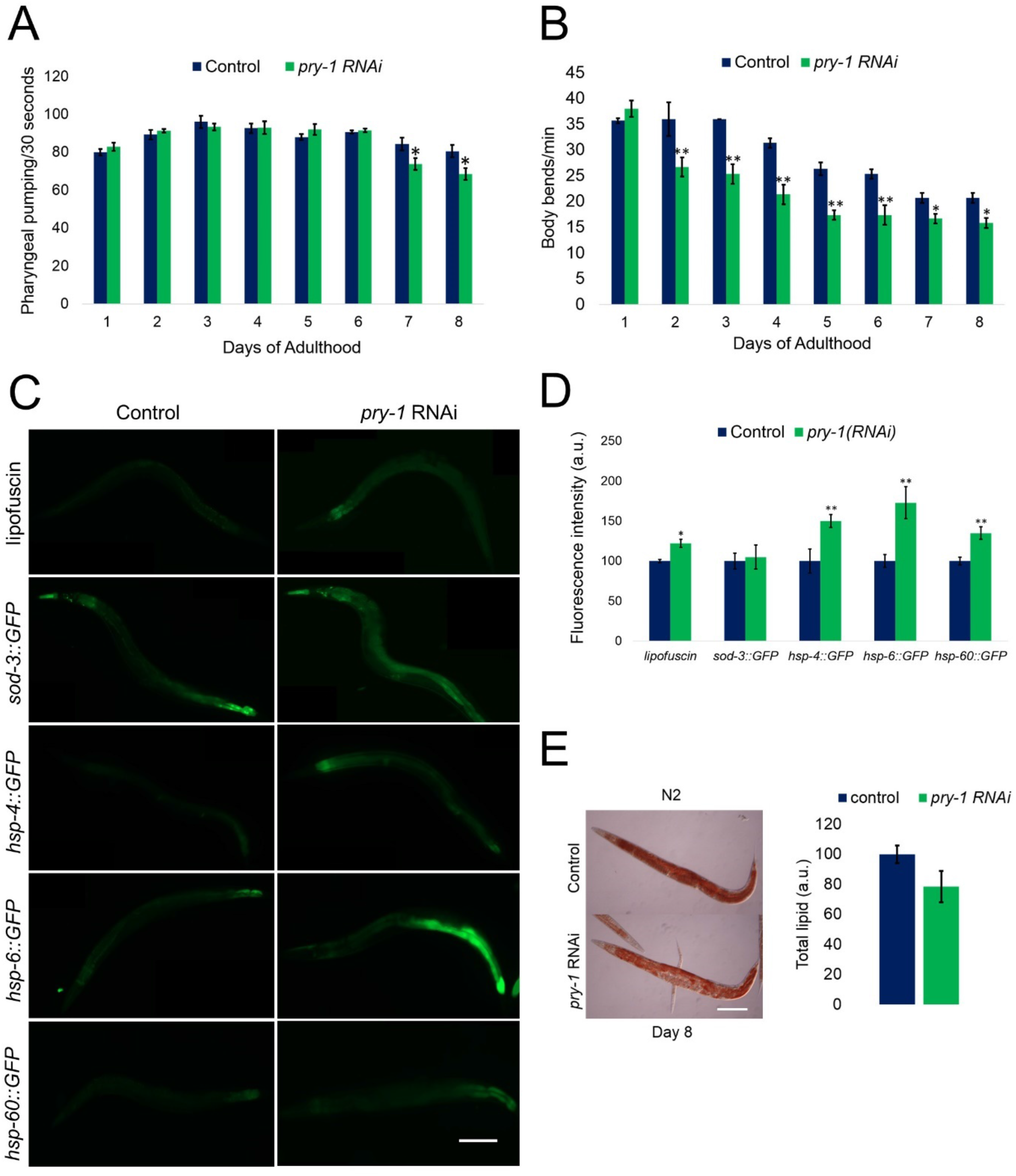
Adult-specific lowering of *pry-1* function accelerates age-associated markers. (A and B) Analysis of the rate of pharyngeal pumping and body bending following *pry-1* RNAi. (A) Animals after *pry-1* RNAi show significantly reduced rate of pharyngeal pumping on day-7 and −8. (B) *pry-1* RNAi treated animals show slower rate of body bending starting day-2 of adulthood. (A and B) Data represents the mean of at least two replicates (at least 30 animals) and error bar represents the standard deviation. Significance was calculated using Student’s t-test \**p* < 0.05, \*\**p* < 0.01. (C) Representative images of animals showing aging pigment (lipofuscin), ROS marker (*sod-3∷GFP*), UPR-ER marker (*hsp-4∷GFP*), and UPR-MT markers (*hsp-6∷GFP* and *hsp-60∷GFP*). Scale bar represents 0.1mm. (D) Quantification of fluorescence intensity shown in figure panel (C). (E) Oil red O (ORO) staining of total lipid droplets in day-8 control and *pry-1(RNAi)* animals. Scale bar represents 0.1mm. (D and E) Data represents the mean of two replicates (at least 15 animals) and error bar represents the standard deviation. Significance was calculated using Student’s t-test. \*\**p* < 0.01.

Next, we measured lipofuscin levels in adults. In *C. elegans*, lipofuscin, a product of oxidative damage and autophagy, is visible as autofluorescent granules in the intestine and serves as a biomarker of aging (Davis et al., 1982). Quantification of the intestinal autofluorescence showed a 1.3-fold increase in *pry-1(RNAi)* adults compared with N2 control animals (**Figure 2C and 2D**). The expression of an oxidative stress marker, manganese superoxide dismutase (*sod-3*), was also investigated (López-Otín et al., 2013). The RNAi-mediated knockdown of *pry-1* caused no significant change in *sod-3*∷*GFP* fluorescence, (**Figure 2C and 2D**), suggesting that *pry-1* function is not essential for the maintenance of oxidative stress. Another indicator of premature aging is the endoplasmic reticulum mediated unfolded protein response (UPR^ER^). Upon activation, UPR^ER^ increases the expression of ER-specific chaperone and heat-shock protein *hsp-4/Bip* (Ron and Walter, 2007). There was a significant increase in *hsp-4∷GFP* expression (1.5-fold) following *pry-1* RNAi-mediated knockdown in day-8 adults (**Figure 2C and 2D**). This observation is supported by the presence of genes in *pry-1* transcriptome involved in IRE-1/IRE1 and PEK-1/PERK-mediated UPR^ER^ signaling (57 genes, 49% overlap, R.F. 3.2, *p*<5.87e-18 and 10 genes, 43% overlap, R.F. 2.9, *p*<0.001; respectively) (**Table S4**), including the key downstream factor XBP-1 that activates *hsp-4* expression (Ron and Walter, 2007). Similar to the UPR^ER^, mitochondrial unfolded protein response (UPR^MT^) is also an indicator of aging. We found that GFP fluorescence of two UPR^MT^ markers, *hsp-6*∷*GFP* and *hsp-60*∷*GFP*, (López-Otín et al., 2013), was increased by 1.7-fold and 1.4-fold, respectively, in *pry-1(RNAi)* day-8 adults (**Figure 2C and 2D**).

Collectively, the data described above provide evidence for *pry-1* playing an essential role in the maintenance of age-associated processes and stress response in animals. One possibility may be that *pry-1* affects aging by regulating lipid metabolism. This is supported by our previous results demonstrating that lipid synthesis is compromised in *pry-1* mutant animals (Ranawade et al., 2018). More importantly, adult-specific knockdown of *pry-1* caused a significant reduction in lipid content in day-8 adults (**Figure 2E**). Given that *daf-16* is also necessary for lipid synthesis (Murphy et al., 2003; Ogg et al., 1997; Zhang et al., 2013a) and *pry-1* and *daf-16* transcriptomes contain a common set of lipid synthesis and transport genes (such as *fat-5-7* and *vit-1/3/4/5*) (**Table S3**), it is conceivable that *pry-1* and *daf-16* interact to regulate lipid levels, leading to a normal lifespan of animals.

### *pry-1* knockdown suppresses lifespan extension of *mom-2/WNT* mutants

Which signaling pathway *pry-1* might utilize to affect the aging process? As PRY-1 is an established negative regulator of WNT signaling, we examined its interactions with WNT ligands. Of the five known ligands, loss of function mutations in *mom-2* and *cwn-2* cause an extension of lifespan (Lezzerini and Budovskaya, 2014). When *pry-1* was knocked down in *mom-2(or42)* and *cwn-2(ok895)* backgrounds, lifespan extension was significantly reduced in *mom-2* mutants (13.6% reduction in mean lifespan, **Figure 3A; Table S2**) but remained unchanged in the *cwn-2* animals (**Figure 3B; Table S2**). We also analyzed the requirements of *bar-1*/*β-catenin*, a component of the canonical WNT signaling that plays a role in aging (Xu et al., 2019; Zhang et al., 2018), in the *mom-2*-*pry-1* pathway. Since *pry-1*-mediated WNT signaling negatively regulates *bar-1*, removing *bar-1* function is expected to suppress the phenotype of *pry-1* mutants. However, we observed that the lifespan of *bar-1* null mutants was further shortened by *pry-1* RNAi (**Figure 3C; Table S2**), suggesting that *bar-1* is unlikely to participate in the *pry-1*-mediated aging process. Further support for this model comes from *bar-1* RNAi experiment that failed to suppress the lifespan phenotype of *pry-1(mu38); hs∷pry-1* animals (**Figure 3D; Table S2**). These data suggest that PRY-1 may act downstream of MOM-2-mediated signaling but independently of BAR-1 in lifespan regulation.

**Figure 3:**
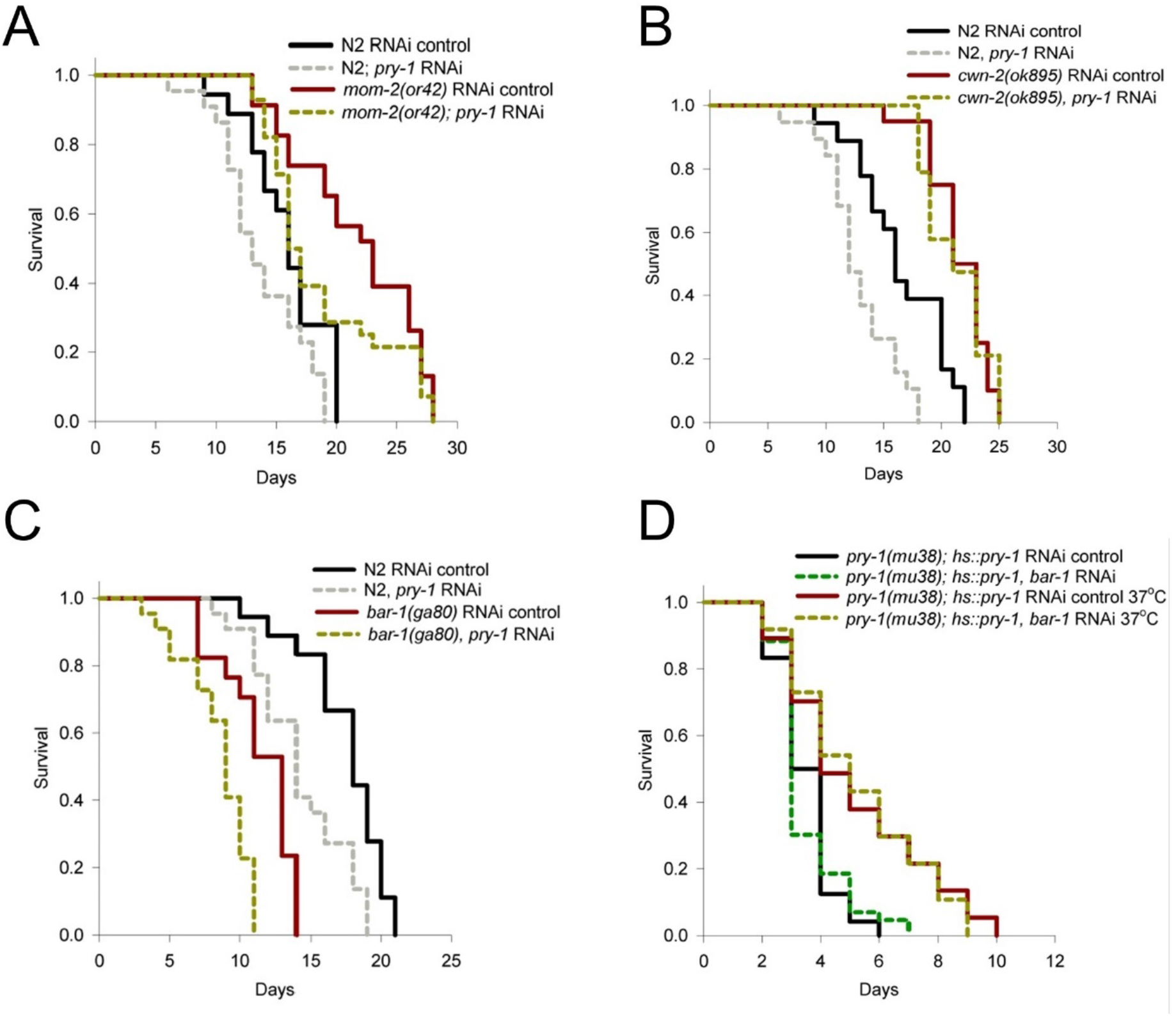
*pry-1* functions downstream of WNT ligand *mom-2* and independently of the β-catenin *bar-1* to regulate lifespan. (A, B and C) Lifespan analysis following knockdown of *pry-1* by RNAi in *mom-2(or42), cwn-2(ok895)* and *bar-1(ga80)* mutants. (A) *pry-1* RNAi reduces lifespan extension seen in the *mom-2* mutants. (B) *cwn-2* mutants show no change in lifespan following *pry-1* RNAi. (C) *pry-1* RNAi exacerbates the shorter lifespan of *bar-1* mutants. (D) *bar-1* RNAi causes no change to the lifespan rescue seen in the *pry-1(mu38); hs∷pry-1* animals. For all the lifespan data with statistics see also STAR methods and Table S2.

### Tissue-specific analysis shows that *pry-1* is needed in muscles and hypodermis

To investigate the requirements of *pry-1* in lifespan regulation, we examined its expression at different stages of life. Previously, a 3.6 kb *pry-1* proximal promoter was used to drive *GFP* reporter, which showed fluorescence throughout development, specifically in the vulval precursor cells, neurons, BWM, and some hypodermal cells (Korswagen et al., 2002). We generated a new *pry-1p*∷*pry-1∷GFP* transgenic strain to characterize expression in detail. The animals showed GFP fluorescence during development in almost all tissues. Expression in seam cells, neuronal cells, muscles, hypodermis, and intestine was readily visible (**Figure 4A**). This pattern of localization matches well with tissue-enrichment of DE genes in the *pry-1* transcriptome (**Table S5**).

**Figure 4:**
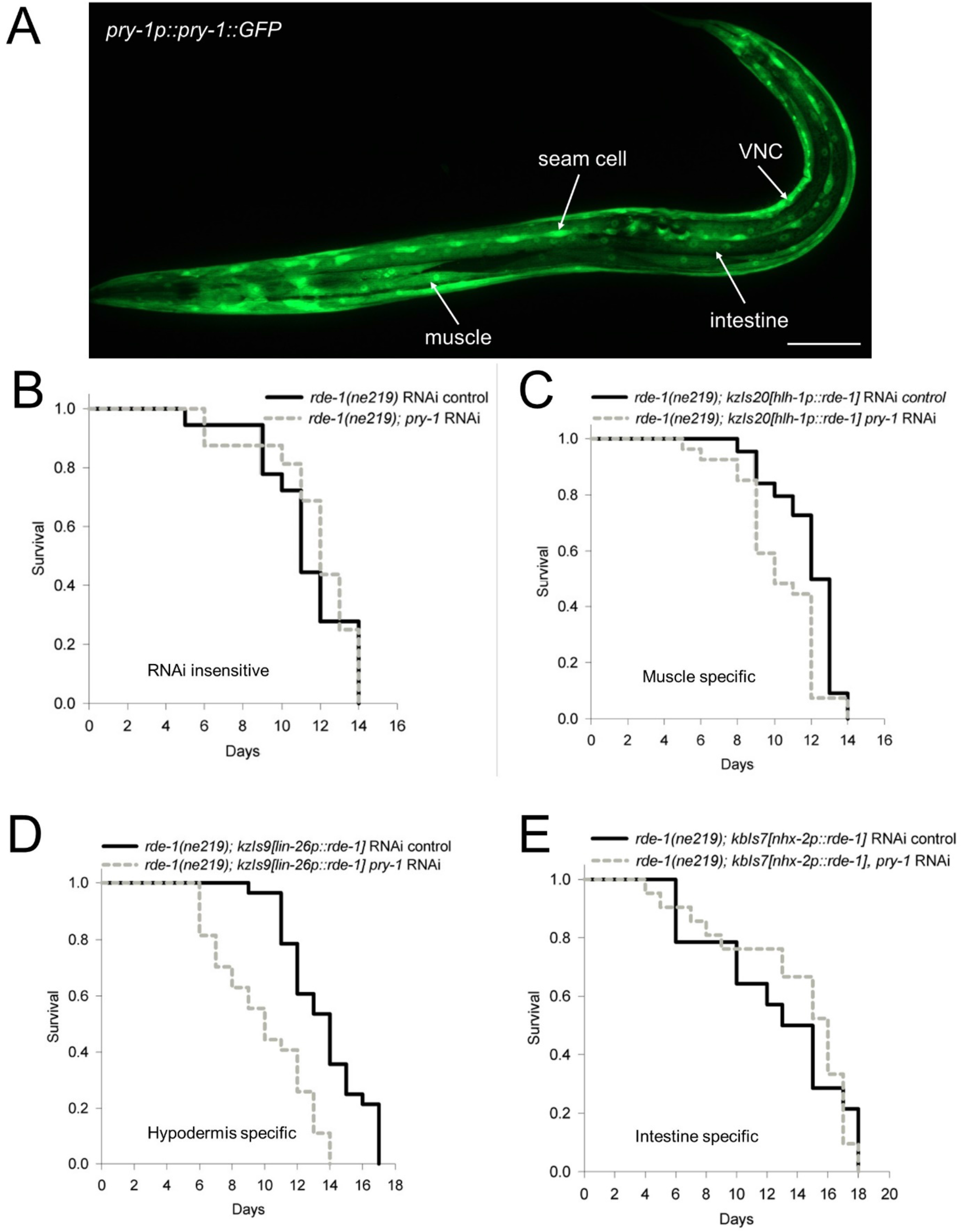
*pry-1* is expressed in multiple tissues during adulthood. *pry-1* function in the muscle and hypodermis is necessary for the normal lifespan. (A) Representative image of *pry-1p∷pry-1∷GFP* animals with expression in the muscle, intestine, seam cell and neurons. Also see Figure S3. Scale bar represents 0.1mm. (B, C, D and E) Lifespan analysis after tissue-specific knockdown of *pry-1.* Also see Figures S4B and S4C. (B) *pry-1* RNAi causes no change in the lifespan of RNAi insensitive animals (*rde-1* mutants). (C and D) Animals have reduced lifespan following *pry-1* knockdown in the muscle and hypodermis. (E) *pry-1* knockdown in the intestine is not detrimental to lifespan. For all the lifespan data with statistics see also STAR methods and Table S2.

A closer examination of GFP analysis in developing animals revealed bright fluorescence in the ventral cord region that includes neuronal and nonneuronal cells. The expression was largely similar in adults although the fluorescence was much brighter in BWM (**Figures 4A, S3A, and S3B**). The posterior end of the intestine, near the rectal opening, showed a strong signal in L4 and adult animals, however, the rest of the intestine lacked a detectable expression. In general, GFP was diffused and not localized to any specific subcellular structures except in the case of muscles and posterior intestine, where nuclei are visible (see arrows in **Figures S3B and S3C**). The fluorescence continued to persist in older adults, consistent with the role of *pry-1* in aging. A similar pattern of expression for *pry-1* was also observed in *C. briggsae* transgenic animals with a marked increase in fluorescence in muscles throughout adulthood (**Figure S4A**). A high level of *pry-1* expression in muscles in both nematodes indicates a conserved role of *pry-1* in maintaining muscle health during aging.

Given that *pry-1* is expressed in muscles as well as other tissues, we examined its tissue-specific requirements for lifespan maintenance. To this end, RNAi experiments were performed in adults using strains that allow tissue-specific knockdowns in muscles, hypodermis, intestine, and neurons (see Star Methods). The results showed that *pry-1* RNAi caused a significant reduction in mean lifespan when knocked down in the hypodermis and muscles (26% lower mean lifespan in hypodermis RNAi and 12% in muscle RNAi) (**Figures 4B-D**). No such effect was observed in other tissues (**Figures 4E, S4B, and S4C**). We conclude that *pry-1* functions in muscles and hypodermis to maintain the lifespan of animals. Further support to this inference comes from the analysis of transgenic strains in which *pry-1* expression was driven by hypodermal and muscle-specific promoters (*lin-26p∷pry-1* and *unc-54p∷pry-1,* respectively). In both cases, the lifespan defect of *pry-1(mu38)* animals was significantly rescued (41% and 56% increase in mean lifespan, respectively by *lin-26p∷pry-1* and *unc-54p∷pry-1*) (**Figures S5A and S5B; Table S2**).

Having uncovered the role of *pry-1* in hypodermis and muscles, we examined whether overexpression of the gene in these two tissues can extend the lifespan. Interestingly, while muscle-specific expression of *pry-1* (*unc-54p∷pry-1*) extended the lifespan significantly (13% increase in mean lifespan), no such effect was observed in the case of hypodermis-specific expression (*lin-26p∷pry-1*) (**Figures 5A and 5B; Table S2**). In fact, *lin-26p∷pry-1* animals were short-lived suggesting that a lack of spatiotemporal control is detrimental (**Figure 5B; Table S2**). These data, together with RNAi and rescue experiments, firmly establish that *pry-1* functions in both muscles and hypodermis for normal lifespan maintenance and that *pry-1* role in hypodermis needs to be tightly regulated. Furthermore, our data has revealed a novel role of *pry-1* in muscles that is beneficial to animals throughout the lifespan. Interestingly, muscle-specific expression of *mAxin1* also caused animals to live longer (14% increase in mean lifespan, **Figure S5C; Table S2**), leading us to propose that Axin family members play similar roles in metazoans.

**Figure 5:**
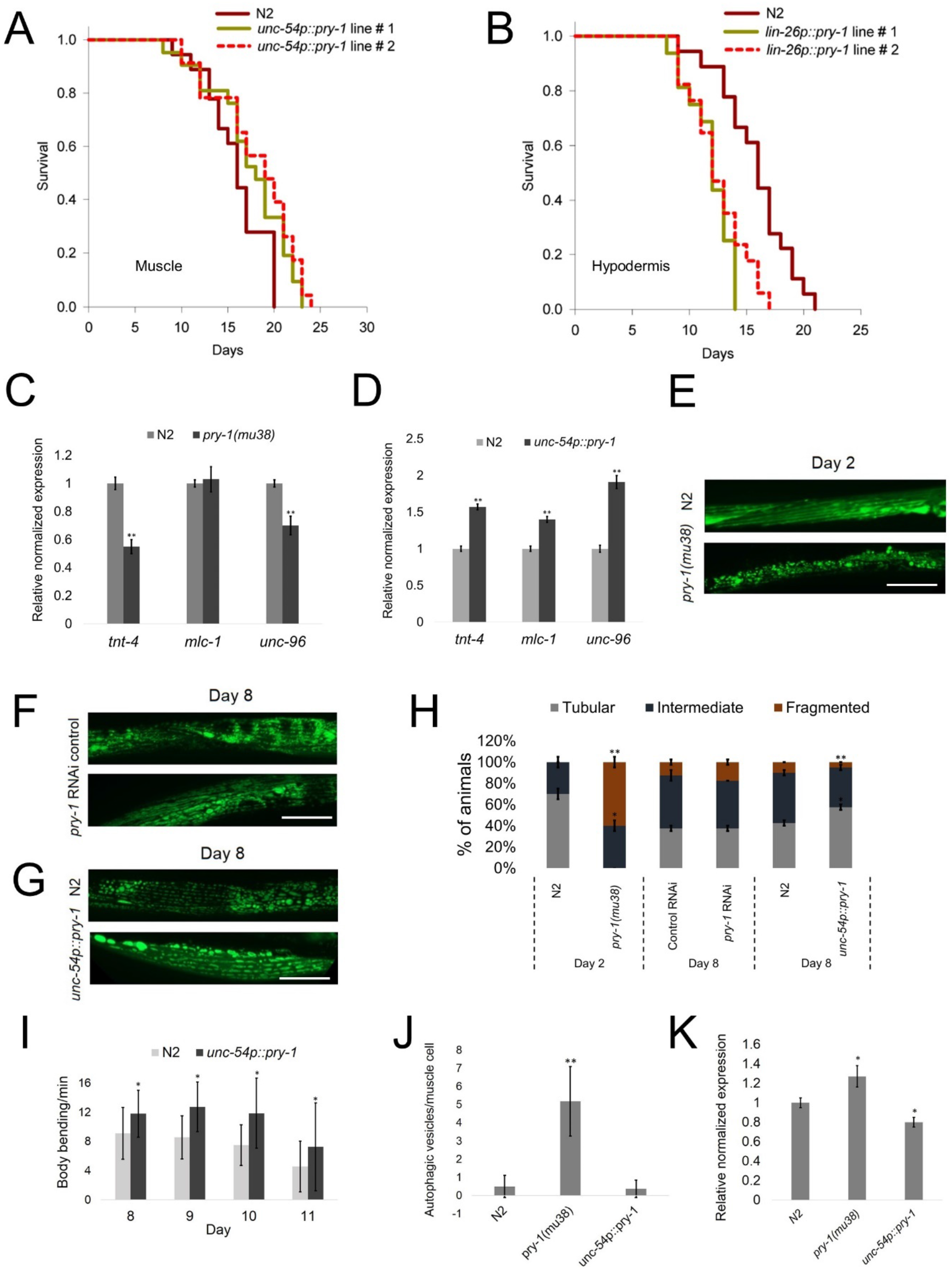
*pry-1* overexpression in the muscle extends lifespan and improves muscle physiology. (A) muscle-specific overexpression of *pry-1* extends lifespan. Also see Figure S5B and Table S2. (B) *pry-1* overexpression in the hypodermis is detrimental to animals. (C and D) qPCR analysis of muscle genes *tnt-4, mlc-1* and *unc-96* in the day-1 *pry-1(mu38)* and *unc-54p∷pry-1* animals. (C) *tnt-4* and *unc-96* are significantly downregulated in the *pry-1* mutants. (D) All three muscle genes are upregulated in the *unc-54p∷pry-1* animals. Data represents the mean of two replicates and error bar represents the SEM. Significance was calculated using Bio-Rad software (Mann–Whitney test). \*\**p* < 0.01. (E, F and G) Representative images for muscle mitochondria morphology in the *pry-1* mutants, *pry-1* RNAi and *unc-54p∷pry-1* animals using *myo-3p∷GFP(mito).* Day-2 adults are analyzed for *pry-1(mu38)* whereas day-8 adults for *pry-1* RNAi and *unc-54p∷pry-1* animals. Scale bar represents 25μm. (H) Quantification of animals with the indicated muscle mitochondria phenotypes. (I) Body bending analysis of *unc-54p∷pry-1* animals between day-8 and day-11. Also see Figure S5D. (J) Number of autophagic vesicles per muscle cell at day-2 of adulthood in animals expressing a GFP marker of autophagic vesicles in body-wall muscles (*dyc-1S∷lgg-1∷GFP*). (H-J) Data represents the mean of two replicates (at least 15 animals) and error bar represents the standard deviation. Significance was calculated using Student’s t-test. \**p* < 0.05, \*\**p* < 0.01. (K) Transcript level of *lgg-1* in the *pry-1* mutants and *unc-54p∷pry-1* animals compared to control. Data represents the mean of two replicates and error bar represents the SEM. Significance was calculated using Bio-Rad software (Mann–Whitney test). \**p* < 0.05.

### Overexpression of *pry-1* in muscles improves muscle health and mitochondrial morphology

The lifespan extension observed in *unc-54p∷pry-1* animals led us to investigate the cellular and molecular basis of *pry-1*’s role in muscle health. We found that *pry-1* transcriptome contains a significant number of muscle genes (31 of 123, 25.2%, R.F. 1.7, *p*< 0.002) (**Table S5**). A majority of these genes are downregulated (90.3%, 28 of 31 genes), suggesting that *pry-1* is needed to maintain their expression. The GO-term analysis identified two broad categories, namely muscle structure development (21 genes) and muscle contraction (15 genes) (**Table S5**), both of which include core components of the sarcomere such as the subunits of troponin complex (*tnt-3*, *tnt-4*), twitchin/titin (*unc-22)*, myosin complex (*mlc-1, unc-15*, *unc-54*), and voltage-gated potassium channel (*unc-58*, *unc-103, slo-1*) (**Table S5**). We chose three genes at random to validate changes in their expression by qPCR (*mlc-1* and *tnt-4*, both involved in muscle contraction and structure development and *unc-96* in muscle structure development). The results confirmed that *tnt-4* and *unc-96* were indeed downregulated in *pry-1(mu38)* animals whereas *mlc-1* expression was unchanged (**Figure 5C**). As expected, all three genes were upregulated in *unc-54p∷pry-1* background (**Figure 5D**).

Since muscle health is linked to mitochondrial homeostasis (Gouspillou and Hepple, 2016; Hood et al., 2019; Lamarche et al., 2018; Regmi et al., 2014), we speculated that *pry-1* might play a role in maintaining the expression of mitochondrial genes. Indeed, genes associated with mitochondrial structure and function were over-represented in the *pry-1* transcriptome (173 genes, 27%, R.F. 1.8, *p*<1.691e^−15^) (**Table S6**). These include genes that function in the mitochondrial membrane (52 of 220, 24% overlap, R.F. 1.6, *p*<5.567e^−4^), mitochondrial outer membrane (10 of 30, 33% overlap, R.F. 2.2, *p*<0.01), mitochondrial matrix (37 of 137, 27% overlap, R.F. 1.8, *p*< 2.298e^−4^), mitochondrial gene expression (18 of 53, 34% overlap, R.F. 2.2, *p*<5.146e^−4^), mitochondrial ATP synthesis (10 of 52, 19% overlap, R.F. 1.3, *p*<0.255), mitochondrial apoptotic changes (2 of 7, 29% overlap, R.F. 1.9, *p*<0.287), mitochondrial fission (3 of 7, 43% overlap, R.F. 2.8, *p*<0.075) and mitochondrial fusion (1 of 5, 20% overlap, R.F. 1.3, *p*<0.440) (**See also Table S6**).

Further support to *pry-1*’s role in mitochondrial health comes from direct visualization of mitochondrial morphology in the BWM (Benedetti et al., 2006). As reported previously, animals exhibit an age-dependent progressive fragmentation of muscle mitochondria with a significant decrease in the proportion of animals showing tubular mitochondrial morphology based on the expression of an organelle-specific GFP reporter, mitoGFP (Lamarche et al., 2018; Regmi et al., 2014). This phenotype reflects the loss of muscle function (Lamarche et al., 2018). We found that while *pry-1* RNAi caused a subtle and statistically insignificant defect in mitochondria in older adults, *pry-1(mu38)* animals exhibited a drastic increase in fragmented mitochondria (**Figures 5E, 5F, and 5H**). By contrast, the morphology was better preserved in *unc-54p∷pry-1* adults (**Figures 5G and 5H**), demonstrating that *pry-1* is needed to maintain the muscle mitochondrial homeostasis.

The above results led us to investigate whether the mitochondrial network architecture mirrors the functional state of muscles. Studies have shown that the loss of locomotion and pharyngeal pumping are associated with the fragmented mitochondrial structure in older worms (Lamarche et al., 2018; Regmi et al., 2014). Since a similar correlation is also seen in *pry-1(mu38)* day-1 adults, we wondered whether *unc-54p∷pry-1* animals will appear healthier with respect to these age-related markers. The experiments revealed that while overexpression of *pry-1* in muscles led to a significantly improved body bending rate in adults, pharyngeal pumping and thrashing were comparable to controls (**Figures 5I and S5D-F**). These results are consistent with *pry-1*’s role in maintaining the mitochondrial network, which may contribute to the betterment of muscle health.

Another process that affects muscle aging is autophagy, in which damaged mitochondria are selectively removed (Madeo et al., 2015; Twig and Shirihai, 2011). While autophagy is beneficial for longevity, its effect is detrimental in the presence of increased mitochondrial permeability that triggers mitochondrial fragmentation (Zhou et al., 2019). Since muscle autophagy increases with age (Lamarche et al., 2018), we investigated whether the process is affected in *pry-1* mutants that are short-lived. The analysis of autophagic vesicles, using *dyc-1S∷lgg-1∷GFP* marker (Lamarche et al., 2018), revealed that vesicle number per muscle cell was significantly higher in *pry-1(mu38)* animals compared to controls (**Figure 5J**), Similar results were also obtained by the analysis of *lgg-1* transcripts (**Figure 5K**). As expected, no such effect was found in *unc-54p∷pry-1* genetic background **(Figure 5J and 5K)**. Altogether, our data has uncovered a new role of *pry-1* in regulating muscle mitochondrial morphology to maintain muscle structure and function.

### *daf-16/FOXO* functions downstream of *pry-1* to maintain the lifespan

As described above, we found that *daf-16* is downregulated in the *pry-1* transcriptome. *daf-16* encodes several isoforms, three of which, R13H8.1b, d, and f, (WormBase WA261 release) influence the rate of the aging process (Kwon et al., 2010). To examine whether *pry-1* affects these isoforms, we performed qPCR analysis. In the case of *pry-1(mu38)*, transcripts for R13H8.1b/c (*daf-16a*) and R13H8.1d/f/h/i/k (*daf-16d/f/h/i/k*) were significantly downregulated (**Figure S6A**). An opposite trend was observed in *unc-54p∷pry-1* animals (**Figure S6B**). How might *pry-1* regulate transcription of *daf-16*? Previously, two intestinal GATA transcription factors *elt-2* and *elt-4* were shown to promote *daf-16* transcription, leading to longevity (Bansal et al., 2014). Using qPCR we found that the expression of both *elt-2* and *elt-4* was significantly upregulated in the muscle-specific line (*unc-54p∷pry-1*) (**Figure S6C**). Thus, *pry-1* may utilize the two GATA factors to affect *daf-16* transcription. Whether this potential mechanism involves an autonomous or non-autonomous function of *pry-1* is currently unknown.

To investigate whether the interaction of *pry-1* with *daf-16* is affected by *daf-2*-mediated signaling (IIS), we knocked down *pry-1* in both *daf-2* and *daf-16* mutant backgrounds. While the knockdown caused a reduction in *daf-2(e1370ts)* lifespan (**Figure S6D**), no change was observed in *daf-16(mu86)* animals (**Figure 6A**), suggesting that *pry-1* may act downstream of *daf-2* but upstream of or in parallel to *daf-16*. The following two experiments are most consistent with the possibility of *daf-16* acting downstream of *pry-1*. One, *daf-16* RNAi suppressed the lifespan extension observed in *pry-1(mu38); hsp∷pry-1* animals (**Figure S6E**). And, two, lifespan defect of *pry-1(mu38)* animals is significantly rescued by *daf-16* overexpression (**Figure 6B**).

**Figure 6:**
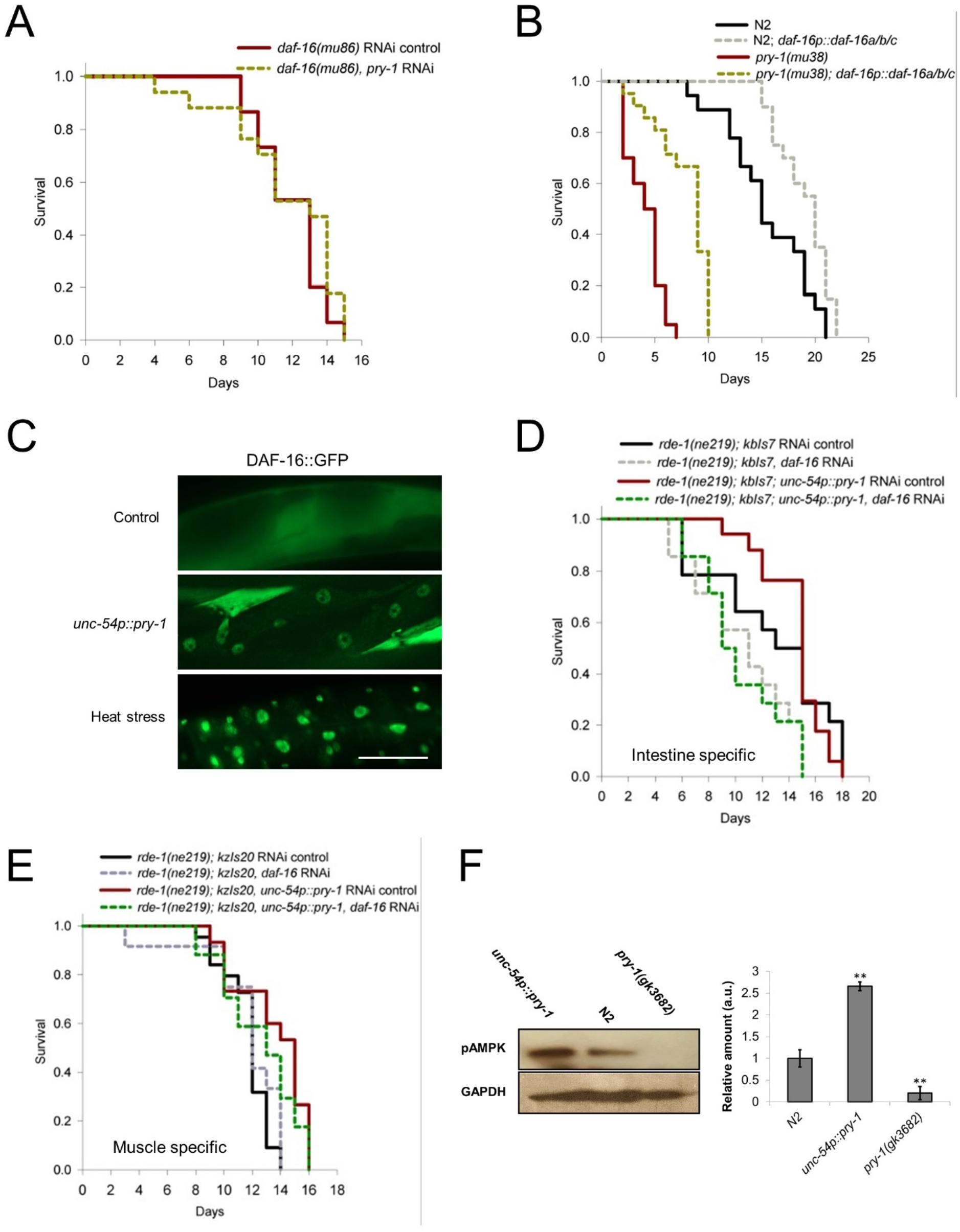
Lifespan regulation by *pry-1* depends on *daf-16* function in the intestine. (A) *pry-1* RNAi does not affect lifespan of *daf-16(mu86)* mutants. (B) *daf-16* overexpression rescues the lifespan defect of *pry-1(mu38)* animals. (C) DAF-16∷GFP is localized in the intestinal nuclei of *unc-54p∷pry-1* animals. Scale bar represents 50μm. (D) Intestine-specific *daf-16* RNAi suppresses lifespan extension of *unc-54p∷pry-1* animals. (E) Muscle-specific *daf-16* knockdown does not affect *unc-54p∷pry-1* animals. (A, B, D and E) For all the lifespan data with statistics see also STAR methods and Table S2. (F) Representative western blotting of AAK-2 phosphorylation in control, *pry-1* mutants and *unc-54p∷pry-1*. Loss of PRY-1 function drastically reduces the amount of activated AAK-2. Data represents the mean of two replicates and error bar represents the standard deviation. Significance was calculated using Student’s t-test. \*\**p* < 0.01.

Since DAF-16’s function depends on its nuclear localization (Libina et al., 2003; Lin et al., 2001), we investigated whether PRY-1 plays a role in this process. The examination of DAF-16∷GFP in *unc-54p∷pry-1* animals revealed that the fluorescence was localized frequently to intestinal nuclei (**Figure 6C**). Consistent with this, *sod-3,* a direct target of *daf-16* was found to be overexpressed (**Figure S6F**). The transgenic worms also exhibited a higher level of lipids (**Figures S7A and S7B**), which along with the known role of *daf-16* in promoting lipid synthesis (Papsdorf and Brunet, 2019) supports the model of *pry-1* interacting with *daf-16* to regulate lipids.

To examine whether *daf-16* acts locally in the intestine or via a long-range signal by functioning in the muscle, we performed tissue-specific RNAi experiments. It was observed that lifespan extension of *unc-54p∷pry-1* was completely abolished by *daf-16* knockdown in the intestine (**Figure 6D**), the tissue where it acts primarily to regulate lifespan (Libina et al., 2003). No such effect was observed following muscle-specific knockdown **(Figure 6E**). Overall, the results show that *daf-16* is involved in *pry-1*-mediated lifespan regulation and that lifespan extension observed in muscle-overexpressed *pry-1* animals depends on nuclear localization of *daf-16* in the intestine.

### DAF-16-mediated PRY-1 signaling depends on AAK-2 function

Having shown that DAF-16 is needed for PRY-1 signaling, we wanted to understand the nature of the interaction between these two factors. In the mammalian system, Axin forms a complex with AMPK upon glucose starvation resulting in phosphorylation of AMPK (Zhang et al., 2013b). An activated AMPK, in turn, phosphorylates a number of targets including FOXO family members, preferentially FOXO3 (Greer et al., 2007a; Mihaylova and Shaw, 2011). Since a similar mechanism was also demonstrated in *C. elegans*, where AAK-2 phosphorylates DAF-16, resulting in DAF-16 nuclear localization and lifespan extension (Greer et al., 2007b; Mair et al., 2011), we investigated whether PRY-1 plays a role in activating AAK-2. For this, AAK-2 phosphorylation was quantified in worm protein extracts. The results showed that while *pry-1* mutants had a drastic reduction in AAK-2 levels when probed with phospho-AMPK antibody, there was a significant increase in AAK-2 in *unc-54p∷pry-1* animals (**Figure 6F**). We also examined whether a drastic reduction in phosphorylated AAK-2 in the *pry-1* mutant could be caused by a lower abundance of AAK-2. The analysis of *aak-2p∷aak-2∷GFP* transgenic animals revealed no change in the fluorescence intensity in *pry-1(mu38)* animals compared to the control (**Figure S7C**). These results demonstrate that PRY-1 is needed for AAK-2 activation, likely via protein-protein interaction.

The above data do not rule out the possibility that PRY-1 acts upstream of AAK-2 in a signaling cascade. Thus, two additional experiments were performed. The results revealed that in one case *pry-1* RNAi did not exacerbate the lifespan defect of *aak-2(ok524)* animals (**Figure 7A; Table S2**). In the other, overexpression of a constitutively active form of AMPKα2 (due to increased T172 phosphorylation) in worms that causes a long-lived phenotype (Mair et al., 2011) was unable to rescue the lifespan defect of *pry-1(mu38)* (**Figure 7B; Table S2**). Together with Western blot data, the findings support a model of PRY-1 regulating AAK-2 phosphorylation, likely through direct interaction, to regulate the lifespan. This conclusion is also supported by expression of *aak-2* in the BWM and neurons during adulthood in a pattern that resembles *pry-1* (Lee et al., 2008; Mair et al., 2011) (**Figure S7D**).

**Figure 7:**
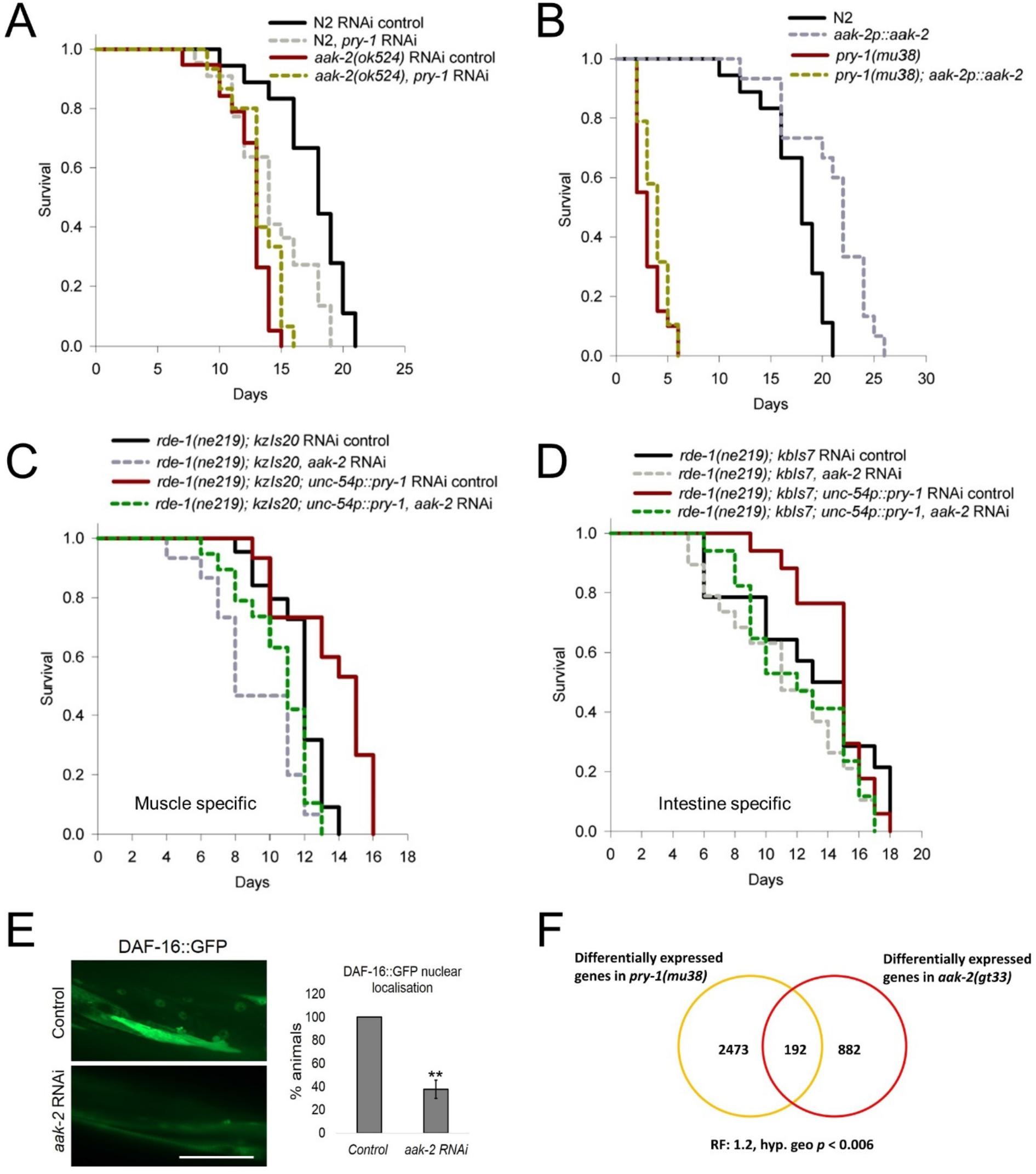
PRY-1 interacts with AAK-2 to regulate DAF-16 localization and extend lifespan. (A) *pry-1* RNAi does not reduce lifespan on *aak-2* mutants. (B) Constitutive activation line of *aak-2* fails to rescue the lifespan defect of pry-1 mutants. (C and D) *aak-2* RNAi in the muscle and intestine suppress the lifespan extension seen in *unc-54p∷pry-1* animals. For all the lifespan data with statistics see also STAR methods and Table S2. (E) *aak-2* RNAi significantly reduces the DAF-16∷GFP nuclear localization seen in the *unc-54p∷pry-1* animals. Data represents the mean of two replicates (30 animals) and error bar represents the standard deviation. Significance was calculated using Student’s t-test. \*\**p* < 0.01. (F) Venn diagram showing an overlapping set of genes between the *pry-1(mu38)* and *aak-2(gt33)* transcriptome.

The above model also suggests that PRY-1 and AAK-2 may regulate a common set of target genes. Indeed, the transcriptome datasets of *pry-1* and *aak-2* mutants (Ranawade et al., 2018; Shin et al., 2011) showed a significant overlap (192 shared genes, 132 up and 60 down; RF: 1.2, hyp.geo *p* < 0.006) (**Figure 7F; Table S7**). Of these, 60 (45%) were found to be mutually upregulated and 28 (47%) mutually downregulated in both the mutants. The overlapping set of DE genes are linked to GO processes such as muscle structure development (*act-1, mel-26, unc-52, emb-9, unc-15* and *unc-54*), muscle contraction (*unc-54*), aging (*daf-16, prmt-1, mpk-1, chc-1, cgh-1, dao-5* and *glp-4*), lipid metabolic process (*tat-4, ldp-1, sptl-3, pmt-1, lipin-1*, and *cgt-3*), and regulation of lipid localization (*daf-16, prmt-1, sams-1, tat-4, lea-1, vit-1, vit-3, vit-4,* and *vit-6*). Moreover, a significant number of genes are associated with stress response (27 genes) and the catabolic process (25 genes) (**Table S7**).

We also determined whether AAK-2 is needed in tissues where PRY-1 and DAF-16 function by performing tissue-specific knockdown experiments. Both muscle and intestine-specific *aak-2* RNAi abolished the lifespan extension of *unc-54p∷pry-1* animals (**Figures 7C and 7D; Table S2**). Also, the RNAi caused significantly fewer animals to show nuclear-localized DAF-16∷GFP (**Figures 7E and F**). Altogether, the data allow us to conclude that AAK-2 interacts with PRY-1 to promote nuclear localization of DAF-16, leading to lifespan extension of animals.

## DISCUSSION

In this paper we report a new role of *C. elegans* Axin family member, PRY-1, in lifespan maintenance that involves the regulation of AAK-2/AMPK-mediated DAF-16/FOXO signaling.

The analysis of *pry-1* transcriptome revealed a significant number of aging-associated genes including IIS and UPR^ER^ pathway components as well as those linked to lipid maintenance. Previous studies have shown that both, DAF-16 and XBP-1-mediated UPR^ER^ signaling, regulate the stress response, lipid metabolism and longevity (Imanikia et al., 2019; Lee et al., 2003; Lin et al., 2018; Murphy et al., 2003; Taylor and Dillin, 2013). We also found that the expression of a significant number of DAF-16 direct targets was altered in *pry-1* mutants and a majority of these were downregulated.

As expected from the misregulation of aging-related genes, a partial or complete loss of PRY-1 activity resulted in the shorter lifespan of animals. The aging phenotype was associated with physiological changes such as slower rates of body bending and pharyngeal pumping, an increase in aging pigment (lipofuscin), and higher expression of UPR^ER^ and UPR^MT^ chaperons. Altogether, these data suggest that *pry-1* affects multiple conserved pathways involved in stress maintenance and aging.

The characterization of *pry-1* expression uncovered muscle as a major site of gene action. Other tissues that express *pry-1* include neurons, hypodermis, and intestine. Since the WNT ligands, *mom-2* and *cwn-2*, are localized in some of these tissues (Song et al., 2010) and both ligands affect lifespan (Lezzerini and Budovskaya, 2014), we investigated the possibility of *pry-1* acting in a WNT-dependent manner. The results of our genetic interaction experiments revealed that *mom-2* and *pry-1* are likely to act in the same pathway to regulate the lifespan in *C. elegans*.

The finding that *pry-1* is expressed in multiple tissues led us to investigate its tissue-specific function. We found that RNAi-mediated knockdowns in muscle and hypodermis caused a significant reduction in lifespan. Moreover, the lifespan defect of *pry-1* mutants was rescued by restoring *pry-1* in muscles and hypodermis. Together, these results demonstrate that *pry-1* is needed in both these tissues to maintain the lifespan. Interestingly, constitutive activation of *pry-1* in muscles but not hypodermis was found to be beneficial to animals. Such a role of *pry-1* does not appear to be unique to *C. elegans* since Axin homologs in other eukaryotes are also expressed in muscles (Smith et al., 2019; Uhlén et al., 2015). Moreover, we found that *mAxin1* was able to extend the lifespan of *C. elegans* when ectopically expressed in the muscle tissue. Thus, the beneficial role of Axin in the muscle may be evolutionarily conserved.

We further explored *pry-1* function in muscle health by analyzing the transcriptome data, which uncovered a significant number of muscle-related genes involved in muscle structure development and function. Almost all of the genes were downregulated. Another group of genes regulated by *pry-1* are associated with mitochondria and include those that function in the mitochondrial membrane, mitochondrial matrix, and mitochondrial ATP synthesis, suggesting that *pry-1* plays a major role in maintaining the health of this vital organelle. As expected, *pry-1* mutants showed increased fragmentation of mitochondria, which appears to contribute to muscle aging and shorter lifespan of the animals (Gouspillou and Hepple, 2016; Hood et al., 2019; Lamarche et al., 2018). Moreover, muscle-specific overexpression of *pry-1* resulted in a marked reduction in the mitochondria phenotype and improved locomotion. Age-associated deterioration in muscle mitochondrial function has been described in other studies. For example, *daf-2* mutants that have a longer lifespan show preservation of mitochondrial morphology and delayed muscle aging (Lamarche et al., 2018; Wang et al., 2019).

To understand how *pry-1* might function in muscle-mediated aging, we investigated the involvement of *daf-16* that plays a role in muscle mitochondrial health (Wang et al., 2019). Our data show that both transcription and subcellular localization of DAF-16 is regulated by PRY-1. Moreover, genetic experiments revealed that the *pry-1*-mediated lifespan depends on *daf-16* and DAF-16 is nuclear-localized when *pry-1* is overexpressed in muscles. Interestingly, the nuclear localization of DAF-16 was observed in the intestine and intestine-specific knockdown of *daf-16* suppressed the lifespan extension of *unc-54p∷pry-1* animals. These findings support a model of *pry-1* activating a long-range signaling to modulate *daf-16* function that in turn may affect cellular processes. One of the outcomes of *pry-1* interaction with *daf-16* could be to affect lipid metabolism, a biological process that is implicated in aging (Papsdorf and Brunet, 2019) since both genes promote MUFA synthesis by transcriptionally regulating fatty acid desaturases such as *fat-7* (Murphy et al., 2003; Ranawade et al., 2018; Zhang et al., 2013a). Other outcomes are also likely since DAF-16 interacts with multiple partners to regulate lifespan (Lapierre and Hansen, 2012; Uno and Nishida, 2016).

Studies have shown that the nuclear localization of DAF-16 depends on its phosphorylation state, a process regulated by the AMPK homolog AAK-2 (Apfeld et al., 2004; Greer et al., 2007b). AMPK-mediated phosphorylation of FOXO has also been demonstrated in the mammalian system (Greer et al., 2007a). We tested whether PRY-1 utilizes AAK-2 to phosphorylate DAF-16, similar to Axin forming a complex with AMPK upon glucose starvation (Zhang et al., 2013b), leading to AMPK activation and phosphorylation of its downstream targets. The results showed that *aak-2* RNAi abolished the nuclear localization of DAF-16 and lifespan extension observed in *unc-54p∷pry-1* animals. Additionally, our Western blot experiments showed that PRY-1 is essential for the activation of AAK-2. Thus, together with experiments involving genetic interactions, AAK-2∷GFP expression, and *pry-1* and *aak-2* transcriptome analysis, the results collectively support the model of PRY-1 and AAK-2 functioning together, potentially forming a complex, to regulate DAF-16 activity for the maintenance of lifespan. It was reported previously that AXL-1, another Axin homolog in *C. elegans*, interacts with AAK-2 following Metformin treatment, although *axl-1* mutants have no age-related phenotypes of their own (Chen et al., 2017). Thus, our work demonstrates that PRY-1 is the sole Axin family member in worms regulating lifespan as well as muscle health.

Interactions between Axin and AMPK have been reported previously in mammalian systems. Specifically, the Axin-AMPK complex formation was enhanced in cultured cells that were subjected to glucose deprivation, and Axin knockdown in the mouse liver impaired AMPK activation (Zhang et al., 2013b). AMPK is known to promote mitochondrial biogenesis and mitochondrial function in human umbilical vein cells and mice aorta (Marin et al., 2017).

Moreover, AMPK phosphorylates all four human FOXO family members (Greer et al., 2007a). Similar to the AMPK, AAK-2-mediated lifespan extension depends on mitochondrial network maintenance and DAF-16 regulation (Greer et al., 2007b; Uno and Nishida, 2016; Weir et al., 2017). Thus, it is plausible that Axin-AMPK-FOXO interact in a conserved manner to regulate disparate biological processes in eukaryotes.

## ACKNOWLEDGMENTS

We thank Wouter Van den Berg, Hannah Hosein and Sakshi Mehta for assistance with experiments. Some of the strains were obtained from CGC, which is funded by the NIH Office of Research Infrastructure Programs (P40OD010440). The tissue-specific promoter plasmids were kindly provided by Richard Roy (McGill University). This work was supported by the NSERC Discovery grant to BG.

## AUTHOR CONTRIBUTIONS

BG conceived and supervised the project. AM, AR, BG designed the study and AM, AR performed the experiments. AM, BG wrote the manuscript and AR contributed to the initial draft. There is no conflict of interest.

## SUPPLEMENTAL INFORMATION

**Figure S1:**
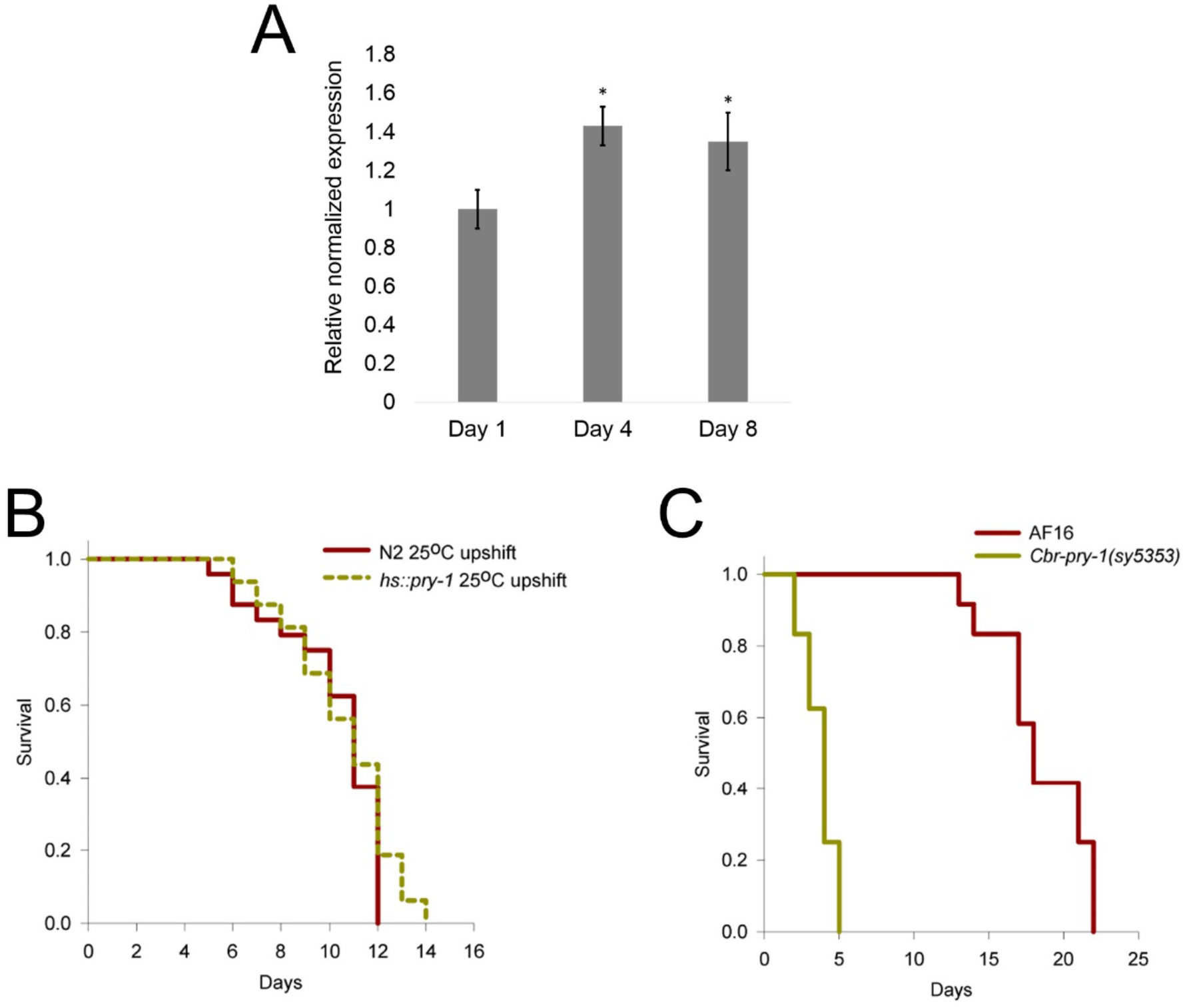
*Cbr-pry-1* mutants have short lifespan. (A) qPCR analysis showing *pry-1* transcript levels at day-1, 4 and 8 of adulthood. Data represents the mean of two replicates and error bar represents the SEM. Significance was calculated using Bio-Rad software (Mann–Whitney test). \**p* < 0.05. (B) Overexpressing *pry-1* during adulthood does not affect lifespan. (C) *Cbr-pry-1(sy5353)* mutants exhibit short lifespan. For all the lifespan data with statistics see also STAR methods and Table S2.

**Figure S2:**
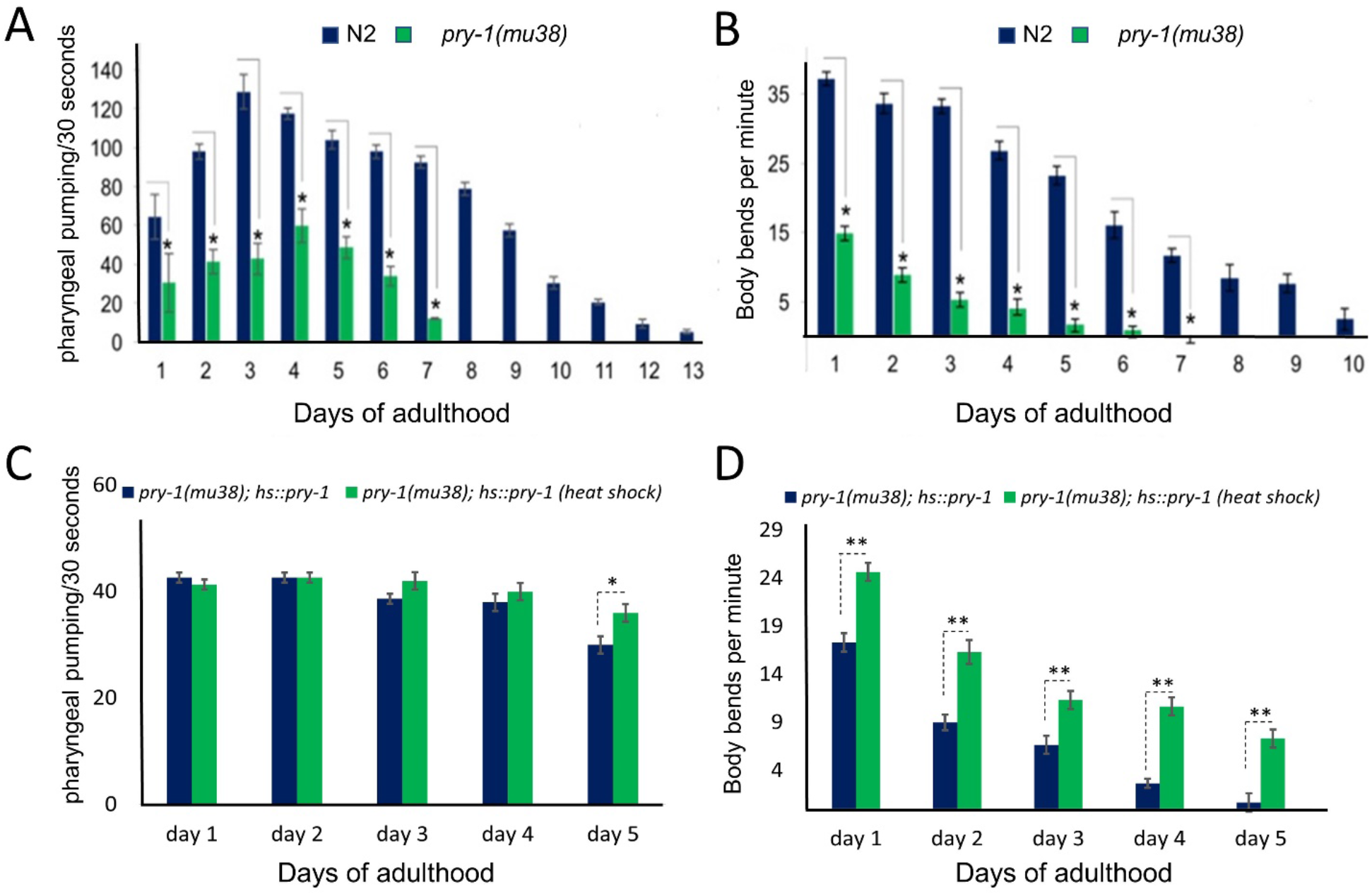
*pry-1(mu38)* mutants show a marked reduction in the rate of pharyngeal pumping and body bending. (A and B) Rate of pharyngeal pumping and body bending in the *pry-1* mutants compared to wild type animals. (C and D) Restoring *pry-1* function in the adult *pry-1(mu38)* animals rescues the defect in the rate of pharyngeal pumping and body bending. Data represents the mean of at least two replicates (at least 30 animals) and error bar represents the standard deviation. Significance was calculated using Student’s t-test \**p* < 0.05, \*\**p* < 0.01.

**Figure S3:**
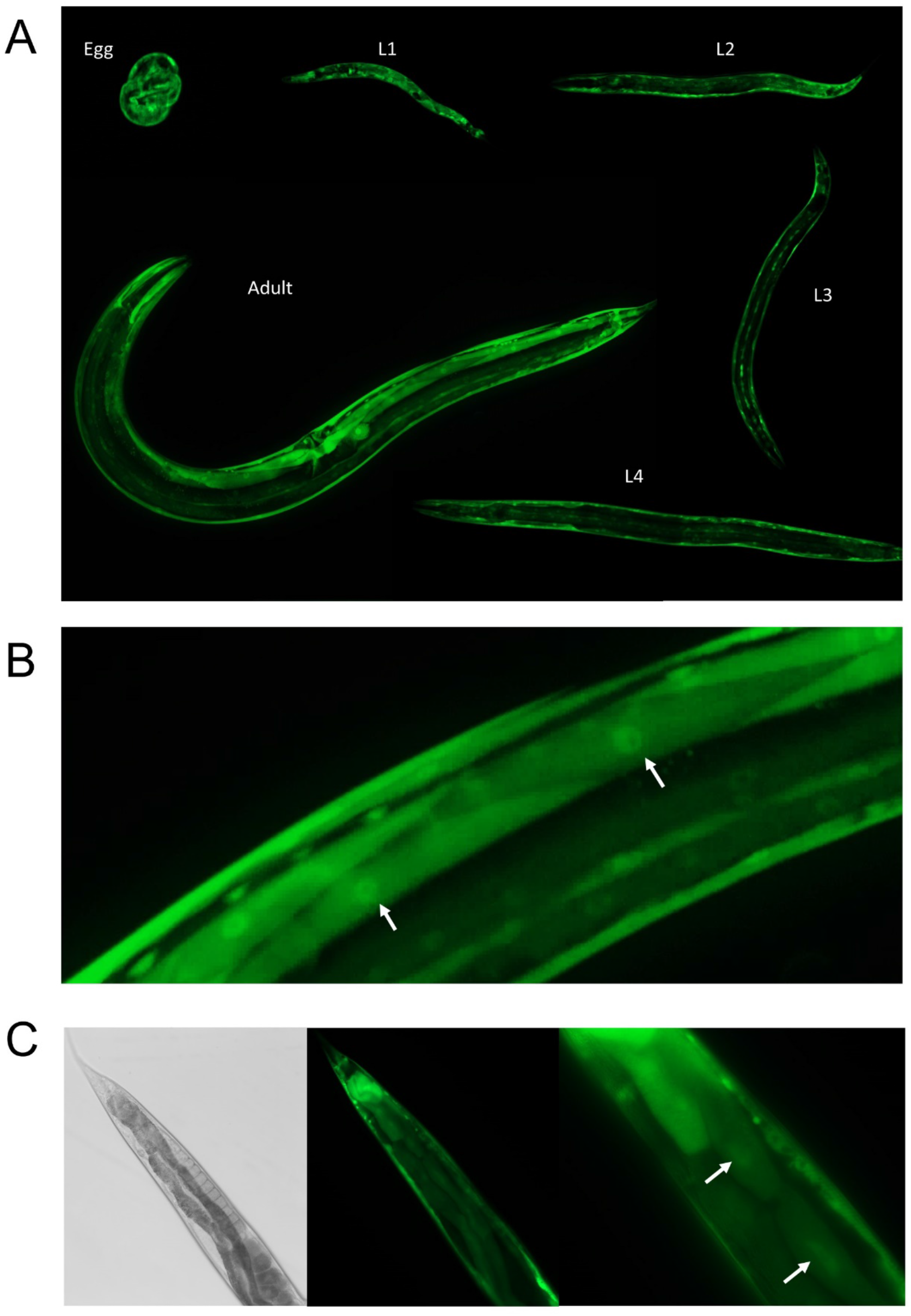
*pry-1* is expressed highly in the muscle throughout lifespan. (A) *pry-1* expression analysis using the *pry-1p∷pry-1∷GFP* transgenic animals at all the larval stages and adults. (B) PRY-1 is found to be diffusely expressed across the body wall muscle cell with higher level in the nucleus (arrowhead). (C) PRY-1 is also present in the intestine (arrowhead showing nucleus).

**Figure S4:**
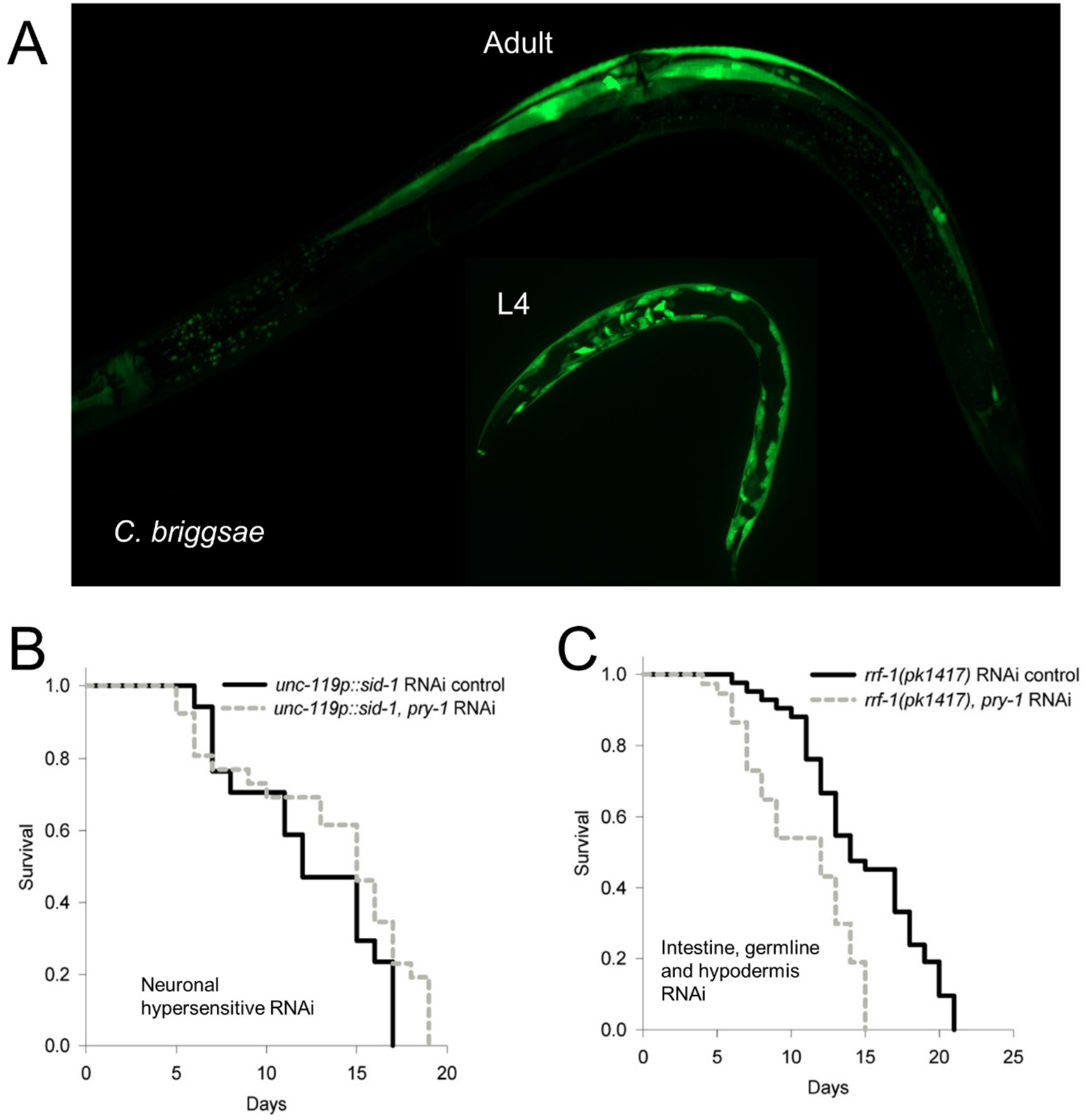
*pry-1* is also expressed in *C. briggsae* muscle tissue. (A) Representative image of *pry-1* expression in the muscle of *C. briggsae* L4 and adult stage animals. (B) *pry-1* RNAi in neurons extends maximum but not mean lifespan. (C) *pry-1* knockdown in the intestine, hypodermis and germline reduces lifespan. For all the lifespan data with statistics see also STAR methods and Table S2.

**Figure S5:**
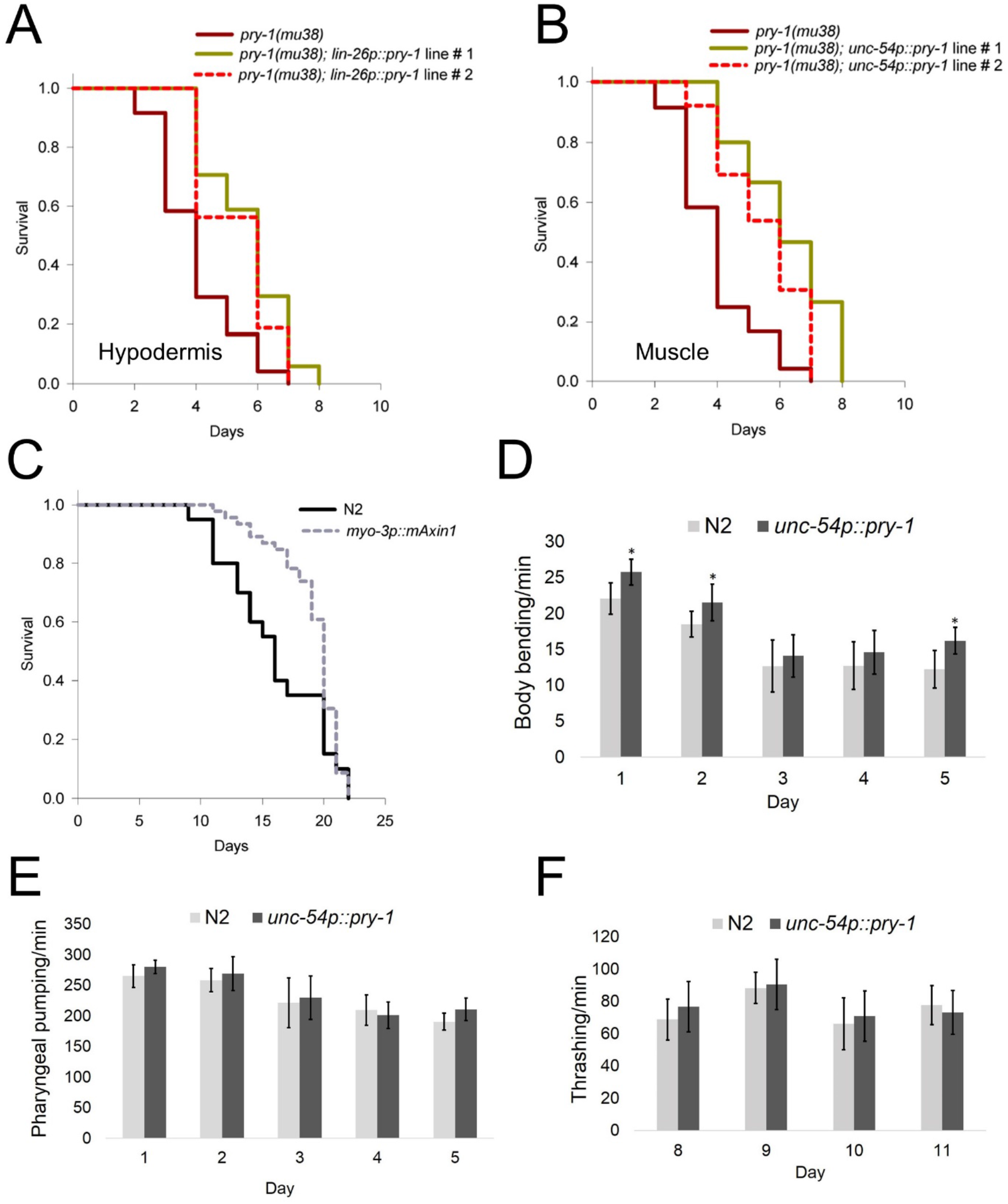
Restoring *pry-1* function in muscle and hypodermis rescues *pry-1(mu38)* lifespan phenotype. (A and B) *pry-1* expression in the hypodermis (*lin-26p∷pry-1*) and muscle (*unc-54p∷pry-1*) rescues *pry-1(mu38)* lifespan. (C) Similar to *C. elegans*, *mAxin1* expression in the muscle extends mean lifespan of animals. For all the lifespan data with statistics see also STAR methods and Table S2. (D and E) Analysis of the rate of body bending and pharyngeal pumping in *unc-54p∷pry-1* animals. (F) Thrashing rate of *unc-54p∷pry-1* between day −8 and −11. (D-F) Data represents the mean of at least two replicates (30 animals) and error bar represents the standard deviation. Significance was calculated using Student’s t-test \**p* < 0.05, \*\**p* < 0.01.

**Figure S6:**
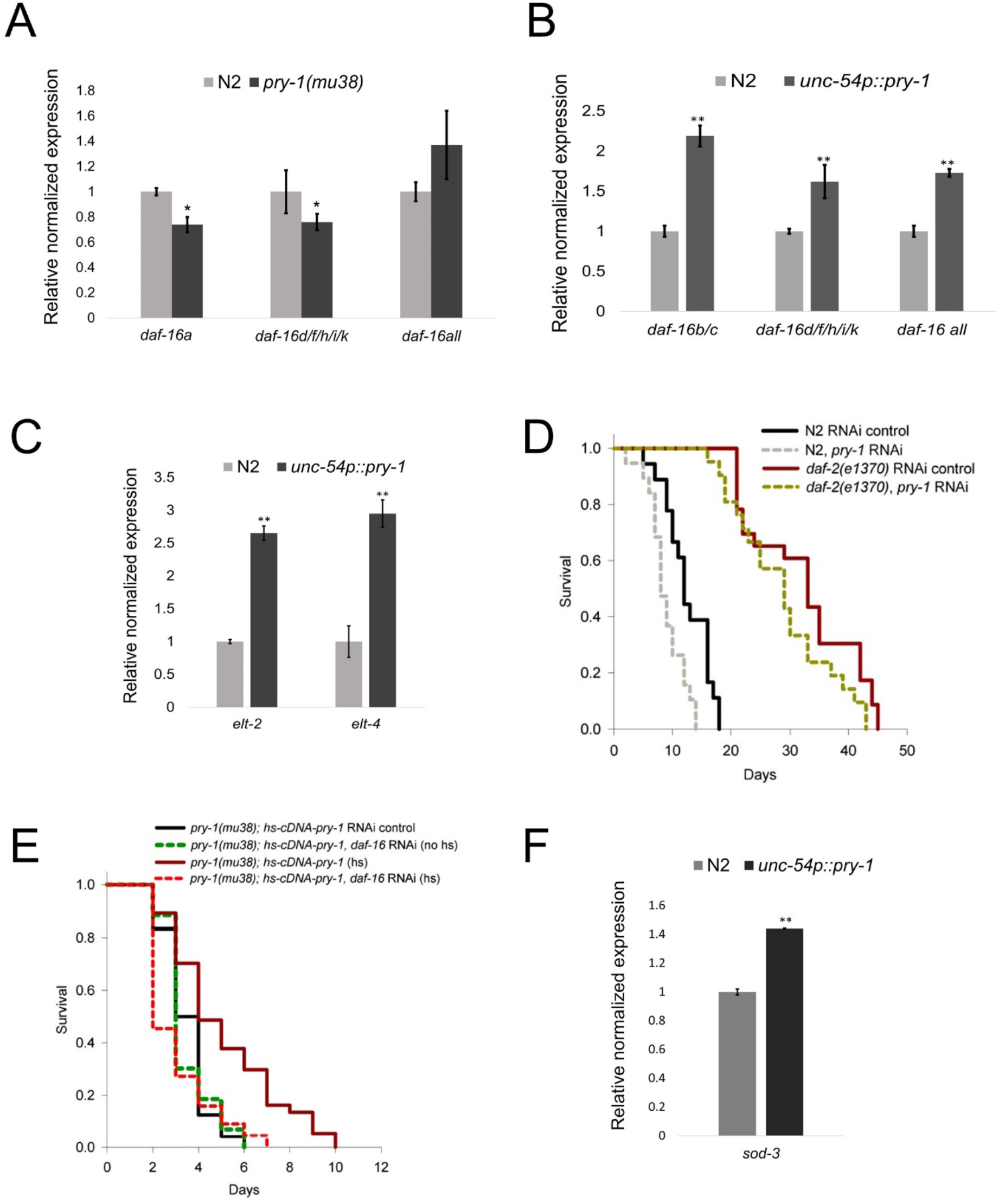
*pry-1* may utilize *elt-2* and *elt-4* to regulate *daf-16* expression. (A and B) Transcript levels of *daf-16a, daf-16d/f/h/i/k* and *daf-16* overall in day-1 *pry-1* mutants and *unc-54p∷pry-1* animals. (A) In the absence of *pry-1* activity, *daf-16* transcripts are significantly down. (B) Overexpression of *pry-1* in the muscle upregulates *daf-16* expression. (C) Transcript levels of *elt-2* and *elt-4* in day-1 *unc-54p∷pry-1* animals. (D) *pry-1* RNAi significantly reduces the longer lifespan of *daf-2* mutants. (E) *daf-16* RNAi suppresses the lifespan rescue seen in the *pry-1(mu38); hs∷pry-1* animals. (D and E) For all the lifespan data with statistics see also STAR methods and Table S2. (F) sod-3 transcript is upregulated in unc-54p∷pry-1 animals. (A, B, C and F) Data represents the mean of two replicates and error bar represents the SEM. Significance was calculated using Bio-Rad software (Mann–Whitney test). \**p* < 0.05, \*\**p* < 0.01.

**Figure S7:**
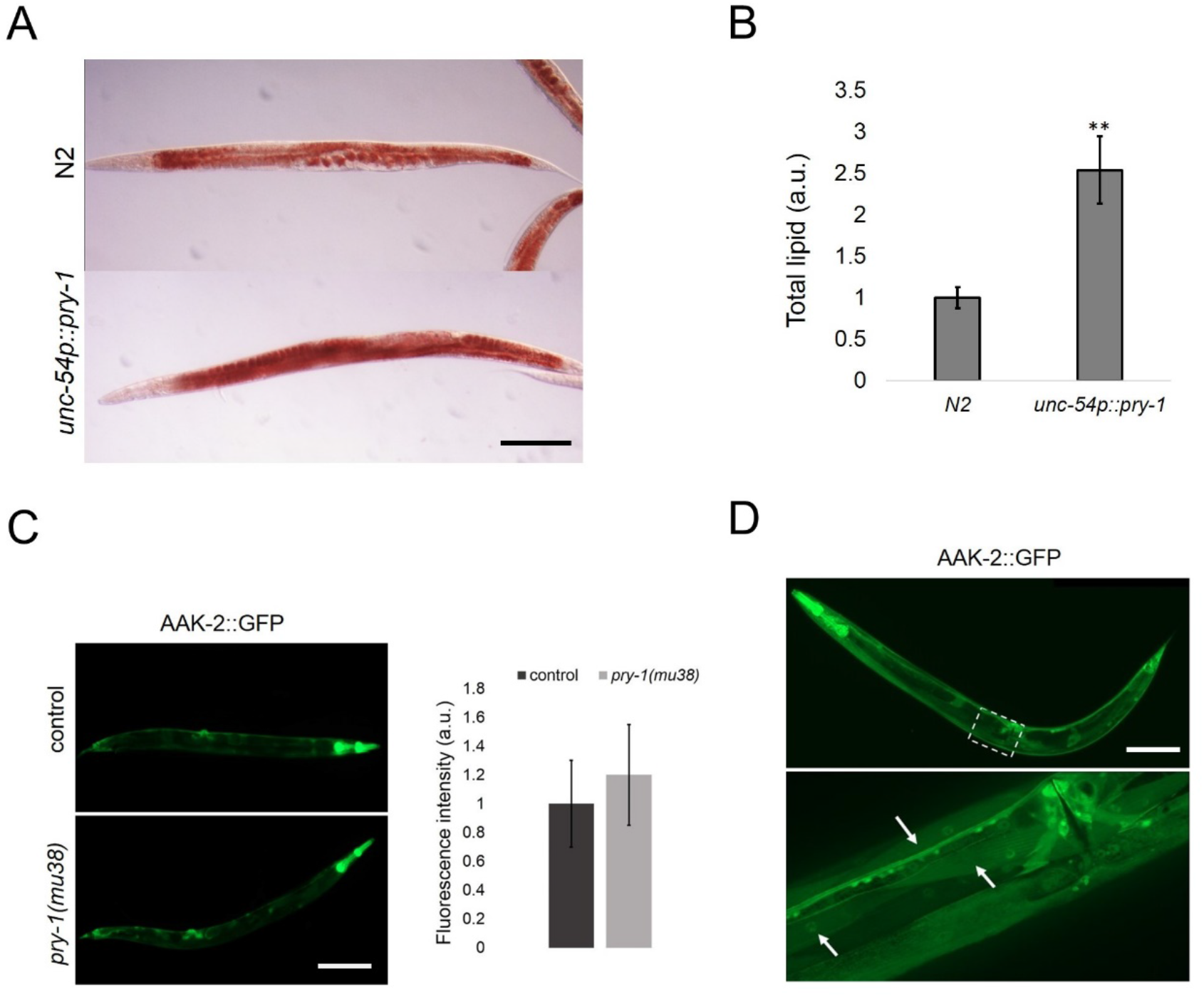
*unc-54p∷pry-1* animals exhibit increased lipid levels. (A) Oil red O (ORO) staining of total lipid droplets in day-1 control and *unc-54p∷pry-1* animals. Scale bar represents 0.1mm. (B) Quantification of lipid levels. Representative image of AAK-2∷GFP in N2 and *pry-1(mu38)* animals. Scare bar represents 0.2mm. Quantification of fluorescence intensity show no change between the two groups. (B-C) Data represents the mean of at least two replicates (at least 20 animals) and error bar represents the standard deviation. Significance was calculated using Student’s t-test, \*\**p* < 0.01. (D) Representative image of AAK-2∷GFP animals. AAK-2 is expressed in the muscle (arrowhead) of adult. Scare bar represents 0.1mm.

## Supplementary Tables

**Table S1: List of aging genes in *pry-1* transcriptomics**.

**Table S2: Aging table showing mean and maximum lifespan of animals. List of primers used in the study**.

**Table S3: Common set of DAF-16 direct targets and DE genes in the DAF-2-DAF-16 pathway**.

**Table S4: DE genes linked to IRE1 and PERK pathways**.

**Table S5: DE genes linked to muscle and Tissue-enrichment analysis (TEA) of DE gene**

**Table S6: DE genes in pry-1 associated with mitochondrial structure and function**.

**Table S7: Common set of DE genes with *aak-2* mutant transcriptomics**.

## REFERENCES

Apfeld, J., O’Connor, G., McDonagh, T., DiStefano, P.S., and Curtis, R.(2004). The AMP-activated protein kinase AAK-2 links energy levels and insulin-like signals to lifespan in C. elegans. Genes Dev. 18, 3004–3009.

Arthur, S.T., and Cooley, I.D. (2012). The effect of physiological stimuli on sarcopenia; Impact of Notch and Wnt signaling on impaired aged skeletal muscle repair. Int. J. Biol. Sci. 8, 731–760.

Bansal, A., Kwon, E.-S., Conte, D., Liu, H., Gilchrist, M.J., MacNeil, L.T., and Tissenbaum, H.A. (2014). Transcriptional regulation of Caenorhabditis elegansFOXO/DAF-16 modulates lifespan. Longev. Heal. 3, 1–15.

Benedetti, C., Haynes, C.M., Yang, Y., Harding, H.P., and Ron, D. (2006). Ubiquitin-like protein 5 positively regulates chaperone gene expression in the mitochondrial unfolded protein response. Genetics 174, 229–239.

Brack, A.S., Conboy, M.J., Roy, S., Lee, M., Kuo, C.J., Keller, C., and Rando, T.A. (2007). Increased Wnt signaling during aging alters muscle stem cell fate and increases fibrosis. Science (80-.). 317, 807–810.

Burkewitz, K., Zhang, Y., and Mair, W.B. (2014). AMPK at the nexus of energetics and aging. Cell Metab. 20, 10–25.

Chen, J., Ou, Y., Li, Y., Hu, S., Shao, L.W., and Liu, Y. (2017). Metformin extends C. Elegans lifespan through lysosomal pathway. Elife 6, 1–17.

Davis, B.O., Anderson, G.L., and Dusenbery, D.B. (1982). Total Luminescence Spectroscopy of Fluorescence Changes during Aging in Caenorhabditis elegans. Biochemistry 21, 4089–4095.

Gouspillou, G., and Hepple, R.T. (2016). Editorial: Mitochondria in Skeletal muscle health, aging and diseases. Front. Physiol. 7, 10–13.

Greer, E.L., Oskoui, P.R., Banko, M.R., Maniar, J.M., Gygi, M.P., Gygi, S.P., and Brunet, A. (2007a). The energy sensor AMP-activated protein kinase directly regulates the mammalian FOXO3 transcription factor. J. Biol. Chem. 282, 30107–30119.

Greer, E.L., Dowlatshahi, D., Banko, M.R., Villen, J., Hoang, K., Blanchard, D., Gygi, S.P., and Brunet, A. (2007b). An AMPK-FOXO Pathway Mediates Longevity Induced by a Novel Method of Dietary Restriction in C. elegans. Curr. Biol. 17, 1646–1656.

Hardie, D.G., and Lin, S. (2019). AMP-activated protein kinase – not just an energy sensor. F1000Research 6, 1–12.

Hardie, D.G., Ross, F.A., and Hawley, S.A. (2012). AMPK: A nutrient and energy sensor that maintains energy homeostasis. Nat. Rev. Mol. Cell Biol. 13, 251–262.

Hood, D.A., Memme, J.M., Oliveira, A.N., and Triolo, M. (2019). Maintenance of Skeletal Muscle Mitochondria in Health, Exercise, and Aging. Annu. Rev. Physiol. 81, 19–41.

Huang, C., Xiong, C., and Kornfeld, K. (2004). Measurements of age-related changes of physiological processes that predict lifespan of Caenorhabditis elegans. Proc. Natl. Acad. Sci. U. S. A. 101, 8084–8089.

Huraskin, D., Eiber, N., Reichel, M., Zidek, L.M., Kravic, B., Bernkopf, D., von Maltzahn, J., Behrens, J., and Hashemolhosseini, S. (2016). Wnt/β-catenin signaling via Axin2 is required for myogenesis and, together with YAP/Taz and tead1, active in IIa/IIx muscle fibers. Dev. 143, 3128–3142.

Imanikia, S., Sheng, M., Castro, C., Griffin, J.L., and Taylor, R.C. (2019). XBP-1 Remodels Lipid Metabolism to Extend Longevity. Cell Rep. 28, 581–589.e4.

Kenyon, C. (2011). The first long-lived mutants: Discovery of the insulin/IGF-1 pathway for ageing. Philos. Trans. R. Soc. B Biol. Sci. 366, 9–16.

Kenyon, C., Chang, J., and Gensch, E. (1993). A C. elegans mutant that lives twice as long as wild type. Nature 366, 461–464.

Klass, M.R. (1983). A method for the isolation of longevity mutants in the nematode Caenorhabditis elegans and initial results. Mech. Ageing Dev. 22, 279–286.

Korswagen, H., Coudreuse, D., Betist, M., Water, S., Zivkovic, D., and Clevers, H. (2002). The Axin-like protein PRY-1 is a negative regulator of a canonical Wnt pathway in C. elegans. Genes Dev. 16, 1291–1302.

Kwon, E.S., Narasimhan, S.D., Yen, K., and Tissenbaum, H.A. (2010). A new DAF-16 isoform regulates longevity. Nature 466, 498–502.

Lamarche, A., Molin, L., Pierson, L., Mariol, M.C., Bessereau, J.L., Gieseler, K., and Solari, F. (2018). UNC-120/SRF independently controls muscle aging and lifespan in Caenorhabditis elegans. Aging Cell 17, 1–11.

Lapierre, L.R., and Hansen, M. (2012). Lessons from C. elegans: Signaling pathways for longevity. Trends Endocrinol. Metab. 23, 637–644.

Lee, H., Jeong, S.C., Lambacher, N., Lee, J., Lee, S.J., Tae, H.L., Gartner, A., and Koo, H.S. (2008). The Caenorhabditis elegans AMP-activated protein kinase AAK-2 is phosphorylated by LKB1 and is required for resistance to oxidative stress and for normal motility and foraging behavior. J. Biol. Chem. 283, 14988–14993.

Lee, S.S., Kennedy, S., Tolonen, A.C., and Ruvkun, G. (2003). DAF-16 target genes that control C. elegans Life-span and metabolism. Science (80-.). 300, 644–647.

De Lencastre, A., Pincus, Z., Zhou, K., Kato, M., Lee, S.S., and Slack, F.J. (2010). MicroRNAs both promote and antagonize longevity in C. elegans. Curr. Biol. 20, 2159–2168.

Lezzerini, M., and Budovskaya, Y. (2014). A dual role of the Wnt signaling pathway during aging in Caenorhabditis elegans. Aging Cell 13, 8–18.

Li, Y.H., and Zhang, G.G. (2016). Towards understanding the lifespan extension by reduced insulin signaling: Bioinformatics analysis of DAF-16/FOXO direct targets in Caenorhabditis elegans. Oncotarget 7, 19185–19192.

Libina, N., Berman, J.R., and Kenyon, C. (2003). Tissue-Specific Activities of C. elegans DAF-16 in the Regulation of Lifespan tissues play an important role in establishing the ani-mal’s rate of aging. First, the C. elegans genome con-tains more than 35 insulin-like genes expressed in a. Cell 115, 489–502.

Lin, K., Hsin, H., Libinia, N., and Kenyon, C. (2001). Regulation of the Caenorhabditis elegans longevity protein DAF-16 by insulin/IGF-1 and germline signaling. Nat. Genet. 28, 139–145.

Lin, X.X., Sen, I., Janssens, G.E., Zhou, X., Fonslow, B.R., Edgar, D., Stroustrup, N., Swoboda, P., Yates, J.R., Ruvkun, G., et al. (2018). DAF-16/FOXO and HLH-30/TFEB function as combinatorial transcription factors to promote stress resistance and longevity. Nat. Commun. 9.

López-Otín, C., Blasco, M.A., Partridge, L., Serrano, M., and Kroemer, G. (2013). The hallmarks of aging. Cell 153, 1194.

Madeo, F., Maiuri, M.C., Kroemer, G., Madeo, F., Zimmermann, A., Maiuri, M.C., and Kroemer, G. (2015). Essential role for autophagy in life span extension. J. Clin. Invest. 125, 85–93.

Mair, W., Morantte, I., Rodrigues, A.P.C., Manning, G., Montminy, M., Shaw, R.J., and Dillin, A. (2011). Lifespan extension induced by AMPK and calcineurin is mediated by CRTC-1 and CREB. Nature 470, 404–408.

Mallick, A., Taylor, S.K.B., Ranawade, A., and Gupta, B.P. (2019a). Axin Family of Scaffolding Proteins in Development: Lessons from C. elegans. J. Dev. Biol. 7, 20.

Mallick, A., Ranawade, A., and Gupta, B.P. (2019b). Role of PRY-1/Axin in heterochronic miRNA-mediated seam cell development. BMC Dev. Biol. 19, 1–12.

Marin, T.L., Gongol, B., Zhang, F., Martin, M., Johnson, D.A., Xiao, H., Wang, Y., Subramaniam, S., Chien, S., and Shyy, J.Y.J. (2017). AMPK promotes mitochondrial biogenesis and function by phosphorylating the epigenetic factors DNMT1, RBBP7, and HAT1. Sci. Signal. 10, 1–12.

Meléndez, A., Tallóczy, Z., Seaman, M., Eskelinen, E.L., Hall, D.H., and Levine, B. (2003). Autophagy genes are essential for dauer development and life-span extension in C. elegans. Science (80-.). 301, 1387–1391.

Mihaylova, M.M., and Shaw, R.J. (2011). The AMPK signalling pathway coordinates cell growth, autophagy and metabolism. Nat. Cell Biol. 13, 1016–1023.

Murphy, C.T., McCarroll, S.A., Lieb, J.D., Bargmann, C.I., Kamath, R.S., Fraser, A., Ahringer, J., Li, H., and Kenyon, C.J. (2003). Genes that act downstream of DAF-16 to influence C. elegans lifespan. Nature 424, 277–283.

Ogg, S., Paradis, S., Gottlieb, S., Patterson, G.I., Lee, L., Tissenbaum, H.A., and Ruvkun, G. (1997). The Fork head transcription factorDAF-16 transduces insulin-likemetabolic and longevity signals in C. elegans. Nature 389.

Papsdorf, K., and Brunet, A. (2019). Linking Lipid Metabolism to Chromatin Regulation in Aging. Trends Cell Biol. 29, 97–116.

Ranawade, A., Mallick, A., and Gupta, B.P. (2018). PRY-1/Axin signaling regulates lipid metabolism in Caenorhabditis elegans. PLoS One 13, e0206540.

Regmi, S.G., Rolland, S.G., and Conradt, B. (2014). Age-dependent changes in mitochondrial morphology and volume are not predictors of lifespan. Aging (Albany. NY). 6, 118–130.

Ron, D., and Walter, P. (2007). Signal integration in the endoplasmic reticulum unfolded protein response. Nat. Rev. Mol. Cell Biol. 8, 519–529.

Seetharaman, A., Cumbo, P., Bojanala, N., and Gupta, B.P. (2010). Conserved mechanism of Wnt signaling function in the specification of vulval precursor fates in C. elegans and C. briggsae. Dev. Biol. 346, 128–139.

Shin, H., Lee, H., Fejes, A.P., Baillie, D.L., Koo, H.S., and Jones, S.J. (2011). Gene expression profiling of oxidative stress response of C. elegans aging defective AMPK mutants using massively parallel transcriptome sequencing. BMC Res. Notes 4.

Smith, C.M., Hayamizu, T.F., Finger, J.H., Bello, S.M., McCright, I.J., Xu, J., Baldarelli, R.M., Beal, J.S., Campbell, J., Corbani, L.E., et al. (2019). The mouse Gene Expression Database (GXD): 2019 update. Nucleic Acids Res. 47, D774–D779.

Song, S., Zhang, B., Sun, H., Li, X., Xiang, Y., Liu, Z., Huang, X., and Ding, M. (2010). A wnt-frz/ror-dsh pathway regulates neurite outgrowth in caenorhabditis elegans. PLoS Genet. 6.

Stenesen, D., Suh, J.M., Seo, J., Yu, K., Lee, K.S., Kim, J.S., Min, K.J., and Graff, J.M. (2013). Adenosine nucleotide biosynthesis and AMPK regulate adult life span and mediate the longevity benefit of caloric restriction in flies. Cell Metab. 17, 101–112.

Taylor, R.C., and Dillin, A. (2013). XXBP-1 Is a cell-nonautonomous regulator of stress resistance and longevity. Cell 153, 1435.

Twig, G., and Shirihai, O.S. (2011). The interplay between mitochondrial dynamics and mitophagy. Antioxidants Redox Signal. 14, 1939–1951.

Uhlén, M., Fagerberg, L., Hallström, B.M., Lindskog, C., Oksvold, P., Mardinoglu, A., Sivertsson, Å., Kampf, C., Sjöstedt, E., Asplund, A., et al. (2015). Tissue-based map of the human proteome. Science (80-.). 347.

Uno, M., and Nishida, E. (2016). Lifespan-regulating genes in c. Elegans. Npj Aging Mech. Dis. 2.

Wang, H., Webster, P., Chen, L., and Fisher, A.L. (2019). Cell-autonomous and non-autonomous roles of daf-16 in muscle function and mitochondrial capacity in aging C. elegans. Aging (Albany. NY). 11, 2295–2311.

Weir, H.J., Yao, P., Huynh, F.K., Escoubas, C.C., Goncalves, R.L., Burkewitz, K., Laboy, R., Hirschey, M.D., and Mair, W.B. (2017). Dietary Restriction and AMPK Increase Lifespan via Mitochondrial Network and Peroxisome Remodeling. Cell Metab. 26, 884–896.e5.

Xu, Y., He, Z., Song, M., Zhou, Y., and Shen, Y. (2019). A micro RNA switch controls dietary restriction induced longevity through Wnt signaling. EMBO Rep. 20, 1–14.

Zhang, P., Judy, M., Lee, S.J., and Kenyon, C. (2013a). Direct and indirect gene regulation by a life-extending foxo protein in C. elegans: Roles for GATA factors and lipid gene regulators. Cell Metab. 17, 85–100.

Zhang, Q., Wu, X., Chen, P., Liu, L., Xin, N., Tian, Y., and Dillin, A. (2018). The Mitochondrial Unfolded Protein Response Is Mediated Cell-Non-autonomously by Retromer-Dependent Wnt Signaling. Cell 174, 870–883.e17.

Zhang, Y.L., Guo, H., Zhang, C.S., Lin, S.Y., Yin, Z., Peng, Y., Luo, H., Shi, Y., Lian, G., Zhang, C., et al. (2013b). AMP as a low-energy charge signal autonomously initiates assembly of axin-ampk-lkb1 complex for AMPK activation. Cell Metab. 18, 546–555.

Zhou, B., Kreuzer, J., Kumsta, C., Wu, L., Kamer, K.J., Cedillo, L., Zhang, Y., Li, S., Kacergis, M.C., Webster, C.M., et al. (2019). Mitochondrial Permeability Uncouples Elevated Autophagy and Lifespan Extension. Cell 177, 299–314.e16.

